# Integrative proteomic and phosphoproteomic profiling of prostate cell lines

**DOI:** 10.1101/696450

**Authors:** Maria Katsogiannou, Jean-Baptiste Boyer, Alberto Valdeolivas, Elisabeth Remy, Laurence Calzone, Stéphane Audebert, Palma Rocchi, Luc Camoin, Anaïs Baudot

## Abstract

**Background:** Prostate cancer is a major public health issue, mainly because patients relapse after androgen deprivation therapy. Proteomic strategies, aiming to reflect the functional activity of cells, are nowadays among the leading approaches to tackle the challenges not only of better diagnosis, but also of unraveling mechanistic details related to disease etiology and progression.

**Methods:** We conducted here a large SILAC-based Mass Spectrometry experiment to map the proteomes and phosphoproteomes of four widely used prostate cell lines, namely PNT1A, LNCaP, DU145 and PC3, representative of different cancerous and hormonal status.

**Results:** We identified more than 3000 proteins and phosphosites, from which we quantified more than 1000 proteins and 500 phosphosites after stringent filtering. Extensive exploration of this proteomics and phosphoproteomics dataset allowed characterizing housekeeping as well as cell-line specific proteins, phosphosites and functional features of each cell line. In addition, by comparing the sensitive and resistant cell lines, we identified protein and phosphosites differentially expressed in the resistance context. Further data integration in a molecular network highlighted the differentially expressed pathways, in particular migration and invasion, RNA splicing, DNA damage repair response and transcription regulation.

**Conclusions:** Overall, this study proposes a valuable resource toward the characterization of proteome and phosphoproteome of four widely used prostate cell lines and reveals candidates to be involved in prostate cancer progression for further experimental validation.

## Introduction

Prostate cancer (PC) is a major public health issue in industrialized countries, mainly because patients relapse by castration-resistant disease after androgen deprivation (1, 2). PC is associated to a panel of clinical states characterized by tumor growth, hormonal status (castration-sensitive or castration-resistant) and presence/absence of metastases. After androgen deprivation therapy, the disease usually progresses to castration-resistant prostate cancer (CRPC), which is highly aggressive and incurable, and jeopardizes the patient’s lifespan and quality of life. This progression involves several molecular mechanisms such as ligand-independent androgen receptor activation or adaptive upregulation of antiapoptotic genes (for review (3)).

Despite an existing treatment guideline for PC and novel clinical trials for CRPC (4, 5), major challenges remain to understand and treat these cancers appropriately. Large-scale *-omics* approaches, able to monitor cancer-induced changes at the cellular level, are among the most promising strategies. Proteomic strategies, by measuring the abundance and activity of proteins, have the ability to directly reflect the functional activity of cells, and to point to deregulations in the most druggable cellular components. In this context, several proteomic studies started to map the landscape of the PC proteome (6–10). These studies identified biomarkers, such as the proneuropeptide *Y*^7^, as well as proteomic changes associated to prostate cancer progression (e.g., increased anabolic processes and oxidative phosphorylation in primary prostate cancer as described by (7) and (8)). Overall, such analyses are valuable not only for diagnosis, but also for providing mechanistic details related to disease etiology and progression.

These proteomic approaches focused on protein quantification, but neglect protein phosphorylation, a key point in the measurement of cellular activity. Protein phosphorylation is a post-translational modification central to signal transduction, that influences cell growth, division, differentiation, cancer development and progression (11, 12). Protein phosphosites can trigger protein activation or inactivation, and profiling the phosphorylation patterns of proteins can be a powerful tool for understanding key roles in tumor progression and/or drug resistance (13). Technological advances in the last decade have led to the development of several highthroughput strategies to map the cellular phosphoproteome (14). Several recent studies examined the phosphoproteome of PC, thereby informing about the activity status of signaling pathways involved in CRPC progression (15–17). In particular, a recent study integrating phosphoproteomics with transcriptomics and genomics data revealed the diversity of activated signaling pathways in metastatic PC patients, in relation to their resistance to the anti-androgen therapy (18). This work further demonstrated the utility of combining *-omics* approaches to better understand PC and CRPC progression. Here, we used a SILAC-based Mass Spectrometry approach, and identified and quantified the proteomes and phosphoproteomes of four widely used prostate cell lines representative of different cancerous and hormonal status. We first identified a common set of housekeeping proteins highly expressed in all cell lines, and enriched in biological processes related to RNA metabolism and oxidative stress. We further detected that each cell line possesses specific protein, phosphosite and functional features, in particular related to cellular metabolism, transport and protein localization. In addition, comparing the sensitive and resistant cell lines, we were able to pinpoint potential biomarkers differentially expressed or phosphorylated in the resistant context. Finally, pathway and network-level interpretation of the biomarkers reveal cellular processes associated with resistance, including, among others, an upregulation of cell migration, extracellular processes and epithelial-mesenchymal transition, and a downregulation of the cellular respiration.

## Methods

### Cell culture and SILAC Labeling

We cultivated three replicates of four cell lines derived from prostate tissue. PNT1A, a non-tumorigenic SV40immortalized human prostatic epithelial cell line (ECACC, European Collection of Cell Cultures, England), castration-sensitive (CS) LNCaP (ATCC, American Type Culture Collection (Rockville, MD, USA)) as well as castration-resistant (CR) DU145 and PC3 cell lines (ATCC). All cell lines were routinely cultured at 37°C in a humidified 5% CO_2_-95% air atmosphere. They were maintained in Dulbecco’s Modified Eagle’s Medium (PC3) and RPMI-1640 (Roswell Park Memorial Institute) (Invitrogen, Cergy Pontoise, France), supplemented with 10% fetal bovine serum (FBS). Stable Isotope Labelling with Amino acids in Culture (SILAC) labeling of cell lines was carried out according to (19, 20) using SILAC media with 10% dialyzed fetal bovine serum supplemented with ^13^C_6_ ^15^N_2_-L-lysine (K8) and ^13^C_6_ ^15^N_4_-L-arginine (R10). Before creating the reference proteome, the incorporation rate of the heavy amino acid was checked for each cell lines using LC-MS/MS and cell extracts were used if this rate reached 95% (data not shown). Additionally, the interconversion of arginine to proline was checked and found to be negligible (data not shown). Cells were washed on ice with PBS and collected in a lysis buffer containing 4% SDS, 100 mM Tris-HCl pH7.4, 1 mM DTT (with protease and phosphatase inhibitors cocktails, EDTA-free, ROCHE, usually 1 tablet of each per 10 ml of lysis buffer). Each pellet was resuspended in the lysis buffer and heated to 95°C for 5 min. Viscous lysates were first homogenized mechanically with a syringe and DNAse was added at a 1:40 dilution (benzonase endonuclease, Sigma). Samples were left on ice for 40 min, then centrifuged at 16000 rcf (g) for 25 min. Supernatants were collected in clean Lo-Bind Eppendorf tubes and protein quantitation was done using BCA assay. After cell lysis, the protein extracts from the four heavy cell lines were mixed in equimolar amounts (1:1:1:1), to generate the super SILAC reference proteome which was then aliquoted and stored at −80°C. For proteomics and phosphoproteomics profiling the reference proteome was mixed in equimolar amounts with protein extracts from each non-labeled cells (Figure 1, C).

**Fig. 1.**
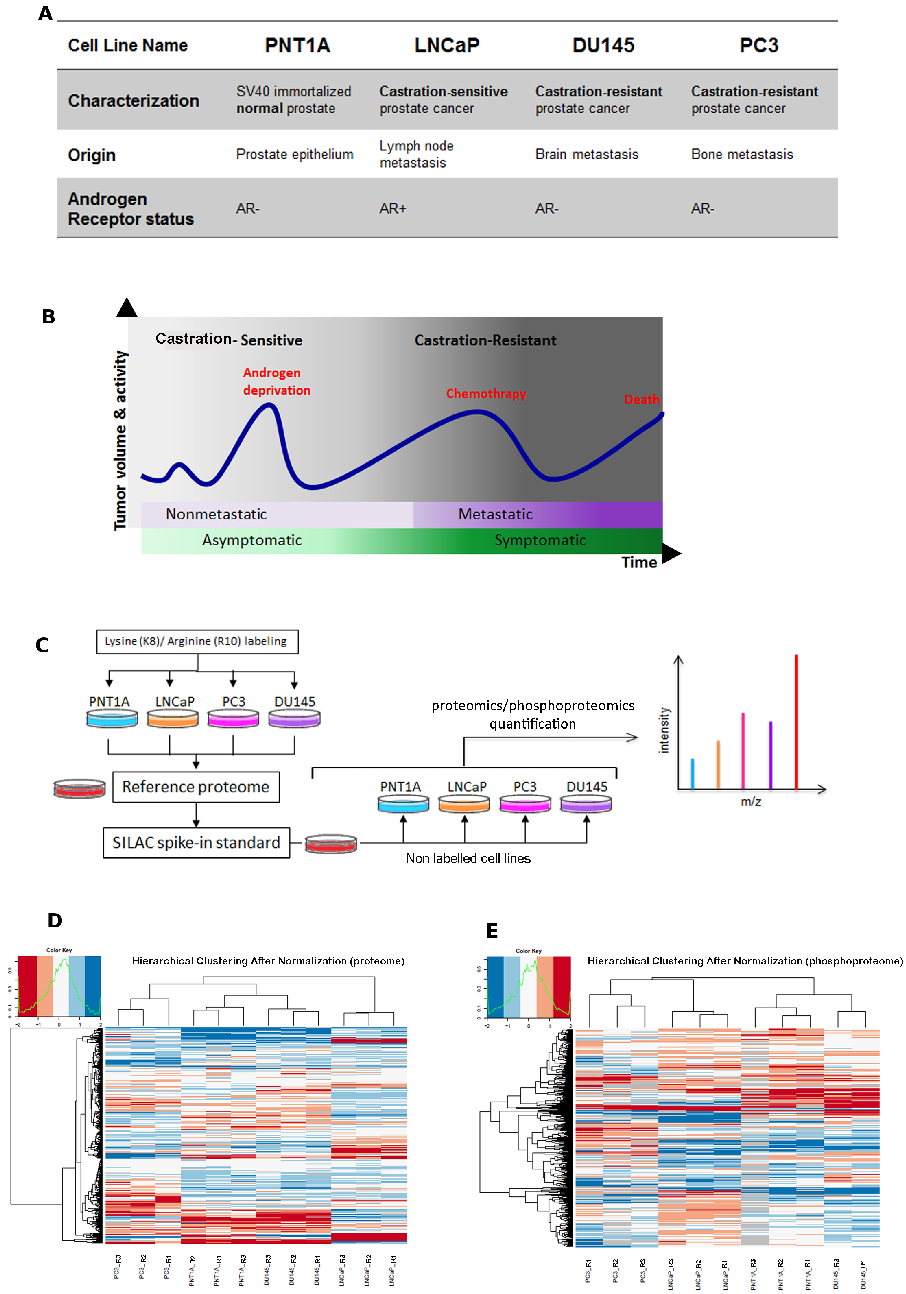
Overall overview of our study. (A) Prostate cell lines used in the present study. AR: Androgen Receptor. (B) Prostate Cancer (PCa) progression over time, from localized asymptomatic castration-sensitive to metastatic castration-resistant disease. (C) SILAC Cell line culture preparation, Spike-in and Mass Spectrometry analysis of the proteomes and phosphoproteomes. Figure adapted from (19) (D) Hierarchical clustering of the proteomes and (E) phosphoproteomes normalized expression data in the four cell lines.

### Proteomes preparation

40 μg of protein extract were loaded on NuPAGE 4–12% bis–Tris acrylamide gels (Life Technologies) to separate proteins, and were stained with Imperial Blue (Pierce, Rockford, IL). Each lane of the gel was cut into 20 bands that were placed in individual Eppendorf tubes. Gel pieces were submitted to an in-gel trypsin digestion using a slightly modified version of the method described by (21). Briefly, gel pieces were washed and destained using few steps of 100mM ammonium bicarbonate. Destained gel pieces were shrunk with 100 mM ammonium bicarbonate in 50% acetonitrile and dried at RT. Protein spots were then rehydrated using 10mM DTT in 25 mM ammonium bicarbonate pH 8.0 for 45 min at 56°C. This solution was replaced by 55 mM iodoacetamide in 25 mM ammonium bicarbonate pH 8.0 and the gel pieces were incubated for 30 min at room temperature in the dark. They were then washed twice in 25 mM ammonium bicarbonate and finally shrunk by incubation for 5 min with 25 mM ammonium bicarbonate in 50% acetonitrile. The resulting alkylated gel pieces were dried at RT. The dried gel pieces were re-swollen by incubation in 25 mM ammonium bicarbonate pH 8.0 supplemented with 12.5 ng/ml trypsin (Promega) for 1h at 4°C and then incubated overnight at 37°C. Peptides were harvested by collecting the initial digestion solution and carrying out two extractions; first in 5% formic acid and then in 5% formic acid in 60% acetonitrile. Pooled extracts were dried down in a centrifugal vacuum system.

### Phosphoproteomes preparation

For each condition, 400 μg of cell lysate implemented with 400 μg of the reference proteome was precipitated using Acetone/Ethanol (sample/Acetone/EtOH 1/4/4 v/v/v) overnight at −20°C. The acetone-precipitated lysate was resolubilized in 50 mM ammonium bicarbonate, pH 8.0. The soluble proteins were reduced for 45 min at 56°C with 10 mM dithiothreitol (DTT), and then alkylated for 30 min at room temperature in the dark with 10mg/ml Iodoacetamide. The protein mixture was then digested with trypsin (1:50 w/w) overnight. Trypsin was quenched by acidification of the reaction mixture with TFA. The peptide mixture was desalted and concentrated on a C18-SepPak cartridge (Waters, Milford, MA) and eluted with 1× 2 mL of 75% acetonitrile (ACN) in 0.1% TFA and dried down. The phosphopeptide enrichment was performed with TiO2 beads 10 μm (Titansphere TIO, GL Sciences, Japan). Titania beads (6 mg) were prepacked in 200 μL pipet tips filled at the orifice with a C8 Empore disk (3M Empore). Prior to loading samples, the titania tips were rinsed with 200 μL of buffer A (3% TFA/70% CH_3_CN). Digest samples were reconstituted with 200 μL of loading buffer (buffer A + 1M Glycolic acid). After centrifugation the supernatant was slowly loaded three times onto the titania tip using centrifugation at 300 g for 10 min. The titania beads were sequentially washed with 200 μL loading buffer, twice with 200 μL of buffer A and 200 μL of 0.1% TFA. Bound peptides were eluted with 140 μL of 1% NH_4_OH pH 10.5 and dried down with a vacuum concentrator.

### Mass Spectrometry analysis

Samples were reconstituted in 0.1% TFA 4% acetonitrile and analyzed by liquid chromatography (LC)–tandem Mass Spectrometry (MS/MS) using Q-Exactive Mass Spectrometer (Thermo Electron, Bremen, Germany) for proteome and phosphopeptide experiments. For the phosphopeptide experiments, an LTQ-Orbitrap Velos Mass Spectrometer (Thermo Electron, Bremen, Germany) was also used. Mass Spectrometers were on line with a nanoLC Ultimate 3000 chromatography system (Dionex, Sunnyvale, CA). Peptides were separated on a Dionex Acclaim PepMap RSLC C18 column at 37°C. First, peptides were concentrated and purified on a precolumn from Dionex (C18 PepMap100, 2 cm × 100 μm I.D, 100 Å pore size, 5 μm particle size) in solution A (0.05% trifluoroacetic acid 2% acetonitrile). In the second step, peptides were separated on a reverse phase column from Dionex (C18 PepMap100, 15 cm × 75 μm I.D, 100 Å pore size, 2 μm particle size) at 300 nL/min flow rate. After column equilibration by 4% of solution B (20% water – 80% acetonitrile – 0.1% formic acid), peptides were eluted from the analytical column by a two steps linear gradient. For proteome analyses, these two steps were 4-25% acetonitrile/H_2_O; 0.1% formic acid for 40 min and 25-50% acetonitrile/H_2_O; 0.1% formic acid for 10 min. For phosphopeptide analyses, these two steps were 4-20% acetonitrile/H_2_O; 0.1% formic acid for 90 min and 20-45% acetonitrile/H_2_O; 0.1% formic acid for 30 min. For peptides ionisation in the nanospray source, spray voltage was set at 1.5 kV and the capillary temperature at 275°C. Instrument method for the Q-Exactive was set up in data dependant mode to switch consistently between MS and MS/MS. MS spectra were acquired with the Orbitrap in the range of m/z 300-1700 at a FWHM resolution of 70000 (AGC target at 1e6, maximum IT 120 ms and 250 ms for proteomes and phosphopeptides respectively). For internal mass calibration the 445.120025 ions was used as lock mass. The 12 most intense ions per survey scan (Intensity threshold 1e5) were selected for HCD fragmentation (AGC target 5e5, NCE 25%, maximum IT 60 ms) and resulting fragments were analysed at a resolution of 17500 in the Orbitrap. Charge state screening was enabled to exclude precursors with unassigned, 1 and > 8 charge states. Fragmented precursor ions were dynamically excluded for 25 s. For phosphopeptides analysis using the LTQ-Orbitrap Velos, the Mass Spectrometer was set as above except for the following parameters. Survey spectra were acquired with a resolution of 60000 (AGC target at 1e6, maximum IT 100 ms) and the 15 most intense precursors ions per cycle were selected for fragmentation by activation of the neutral loss ions (−48.99, −32.66, and −24.49 Thompson relative to the precursor ions) with collision induced dissociation (AGC target 3000, NCE 35%, maximum IT 200 ms). The Mass Spectrometry proteomics data, including search result, have been deposited to the ProteomeXchange Consortium (www.proteomexchange.org)(22) via the PRIDE partner repository with datasets identifiers PXD004970 and PXD004992.

### Protein identification and quantification

Relative intensity-based SILAC quantification was processed using MaxQuant computational proteomics platform, version (23). First the acquired raw LC Orbitrap MS data were processed using the integrated Andromeda search engine (24). Spectra were searched against a SwissProt human database (version 2014.02; 20284 entries). This database was supplemented with a set of 245 frequently observed contaminants. The following parameters were used for searches: (i) trypsin allowing cleavage before proline (25); two missed cleavages were allowed; (ii) monoisotopic precursor tolerance of 20 ppm in the first search used for recalibration, followed by 6 ppm for the main search and 20 ppm for fragment ions from MS/MS; (iii) cysteine carbamidomethylation (+57.02146 Da) as a fixed modification and methionine oxidation (+15.99491 Da) and N-terminal acetylation (+42.0106 Da) as variable modifications; (iv) a maximum of five modifications per peptide allowed; and (v) minimum peptide length was 7 amino acids. The re-quantify option was enabled to search for missing SILAC partners. The quantification was performed using a minimum ratio count of 2 (unique+razor) and the second peptide option to allow identification of two co-fragmented co-eluting peptides with similar masses. The false discovery rate (FDR) at the peptide level and protein level were set to 1% and determined by searching a reverse database. For protein grouping, all proteins that cannot be distinguished based on their identified peptides were assembled into a single entry according to the MaxQuant rules.

### Phosphopeptide identification and quantification

Peptide identification was done similarly than above using MaxQuant software except that serine, threonine, and tyrosine phosphorylation (+79.96633 Da) were allowed as variable modifications.

### Preliminary treatment of the datasets

Statistical analyses were done with the Perseus program (version 1.3.0.5; freely available at www.maxquant.org) from the MaxQuant environment (26). The relative intensity-based SILAC ratio, iBAQ normalised intensities and peptide intensities were uploaded from the proteinGroups.txt and Phospho(STY)Sites.txt files for proteome and phosphoproteome studies, respectively. Proteins marked as contaminant, reverse hits, and “only identified by site” were discarded.

One DU145 cell line replicate in the phosphoproteome study was discarded due to high divergence. In all other cases, for each experiment and for each cell line, the measurements of three replicates were considered. We identified 3219 proteins (FDR 1% for peptide and protein identification) in triplicates (Supplementary Table 1). We kept for further quantification analyses only those proteins containing at least two valid values (over the 3 replicates) in each cell line. This very conservative approach avoids imputing missing values, and ensures the results of the statistical tests. Doing so, we quantified 1229 proteins (Supplementary Table 1), used for all subsequent analyses. For the phosphoproteomics analysis, a similar strategy allowed identifying 3746 phosphosites, of which 563 were kept for further quantification analysis following the previously defined filters (Supplementary Table 2).

### Data analyses

R statistical programming environment (27) was used for the treatment of the proteomic and phosphoproteomic datasets. Expression ratios towards the internal standard were base-2 logarithmized and normalized using z-scores.

#### Clustering

Unsupervised hierarchical clustering using average method was performed for the proteomic and phosphoproteomic datasets based on Euclidean distances of the expression ratio after normalization.

#### Identification of the highly-expressed housekeeping proteome

The abundance of each protein in each cell line was computed as the sum of the IBAQ values of every replicate. The housekeeping proteome was obtained by selecting the 10% most abundant proteins matching across all cell lines.

#### Identification of differentially expressed proteins and phosphosites

We first applied a 1-way ANOVA over the four different cell lines. Benjamini & Hochberg FDR (28) was used for multiple testing corrections, and the threshold for significance was set to 0.01.

Next, to characterize cell line specific protein/phosphosite expression, a t-test was applied to compare the expression value in the three PC cell lines (LNCaP, DU145 and PC3) to the reference non-tumorigenic PNT1A cell line. Benjamini & Hochberg FDR (28) was used for multiple testing corrections, and the threshold of significance set to 0.1.

Pairwise comparisons of protein/phosphosite expression values between the castration-sensitive (CS: LNCaP) and the castration-resistant cell lines (CR: DU145 and PC3) were performed with a t-test, and the threshold of significance set to 0.1 after FDR multiple testing corrections. The results of the pairwise comparisons with the two CR cell lines were combined to define proteins/phosphosites always significantly up- or downregulated in CS as compared to CR.

It is to note that these analyses are conducted with a very stringent filter that select only proteins and phosphosites with at least two over three valid quantification values in all four cell lines. In this context, the proteins identified only in the CR resistant or only in the CS sensitive contexts were discarded, whereas they could be considered as pertinent biomarkers. We then also rescued these potential biomarkers as “CR_only” proteins and phosphosites, having at least two valid expression values in CR and strictly none in CS cell lines and “CS_only” proteins and phosphosites, having at least two valid values in the two CS cell line and strictly none in the CR cell lines.

#### Pathway and biological process analyses

##### Functional Enrichments

Enrichment Analyses were conducted with G:Profiler (29), and the significance threshold set to 0.01 after FDR multiple testing corrections. The list of 1229 proteins used for quantification analyses was used as statistical background. Additionally, the strong filter option was selected on G:Profiler to display solely the most significant ontology in each ontological group, and reduce annotation redundancy.

#### ROMA

ROMA (Representation and Quantification of Module Activity) is a software focused on the quantification and representation of biological module activity using expression data (30). The reference gene sets used for this analysis were selected from pathway databases including Reactome (31) and HALLMARK (32). For each of these pathways, a score corresponding to the weighted sum of the protein expression was computed. The weights are based on the first principal component (PC1). ROMA quantifies the statistical significance of the amount of variance explained by the PC1, and is referred to as the gene set overdispersion. Overdispersed pathways are selected based on a p-value set to 0.01, and the resulting list of pathways can be interpreted as the pathways that contribute significantly to the total expression variance. A detailed presentation of the computational method and use of software can be found at (30). For this study, we applied on the proteomic dataset an R implementation of ROMA (https://github.com/Albluca/rRoma), which is an improved version of the initial software. The results are presented as a heatmap where the mean value of the scores was computed by types of cancer cell lines: CS for castration-sensitive and CR for castration-resistant, and scaled between −1 and 1.

#### Ingenuity Pathway Analysis (IPA)

Proteomic datasets were also analyzed with Ingenuity Pathway Analysis (IPA) software (Qiagen, http://www.ingenuity.com/) to predict pathway activa tion or inhibition. The IPA knowledgebase, derived from literature, compute a score based on one-tailed Fisher test. The final score corresponds to the negative log of p-value, and thresholds were set to 0.01.

#### KSEA

In order to use the KSEA App (https://casecpb.shinyapps.io/ksea/) (33) on the phosphoproteomic datasets, we computed the fold-changes (FC) between DU145 and LNCaP, and between PC3 and LNCaP, using the mean raw expression values of the replicates. We selected the sites where the expression values are over or under-expressed in both CR cell lines in comparison with LNCaP. Finally, we computed the mean of the FC for the 337 Sites, and normalized it between 0 and 1.

We used this list of sites as input for the KSEA App. The kinases with at least 3 targeted phosphosite substrates, and a p-value smaller than 0.05 were considered as significant.

#### Network analyses

We constructed a network encompassing molecular complex interaction data by merging Corum complexes (34) and Hu.MAP complexes (35). This network contains 8653 nodes and 91500 edges. Then, we fetched interactions between:

- Proteins significantly differentially expressed in CR versus CS;
- Proteins containing phosphosites significantly differentially expressed in CR versus CS;
- Proteins and proteins containing phosphosites identified only in CR or CS contexts (CR_only and CS_only).

Overall, the interactions between these proteins led to an interaction network of 359 nodes and 1161 edges, including a large connected component encompassing 194 nodes and 1098 edges represented with Cytoscape (36). For visualization purposes, the expression values mapped on the network in Figure 4 correspond to the mean of the expression of PC3 and DU145 cell lines.

## Results

### Proteomic and phosphoproteomic profiles of prostate cell lines

In order to elucidate prostate cancer progression and androgen escape pathway with proteomics and phosphoproteomics identification and quantification, we selected four widely exploited prostate cell lines, namely PNT1A, LNCaP, DU145 and PC3 for proteomic and phosphoproteomic profiling (Figure 1, A). These cell lines are routinely used, and are representative of normal, cancerous and castration-resistant progression of prostate cancer (Figure 1, B). The PNT1A benign prostate cell line was established by immortalizing non-tumorigenic human prostate benign epithelial cells by transfection with the SV40 large-T antigen gene (37). The castration-sensitive LNCaP cell line was established from metastatic deposit in a lymph node and demonstrates androgen sensitivity (38). Finally, the two castration-resistant tumor cell lines, DU145 and PC3, were established from metastatic deposits (bone/lumbar spine and central nervous system, respectively), lack the androgen receptor (AR) and are androgen-independent. Moreover, PC3 cells are more tumorigenic and have a higher metastatic potential than DU145 (39). It is to note that the benign PNT1A cell line also lacks the AR. The loss of AR and prostate-associated markers (PSA and PAP) appears to be a consistent feature of immortalized cells of prostatic origin, observed in SV40 immortalized cell lines such as PNT1A (40).

We used Stable Isotope Labelling with Amino acids in Culture (SILAC) and Mass Spectrometry (MS) to identify and quantify the proteomes of these four cell lines (41, 42) (Figure 1, C, Methods). We elected the spike-in super SILAC method described by (19, 20). In this protocol, the protein expression in each cell line is compared to the same reference proteome, thereby maximizing the number of detected proteins. We identified 3219 proteins (Supplementary Table 1). We plotted the median iBAQ values considering all the cell lines to estimate the absolute abundance of the 3219 identified proteins, and obtained the expected S-shaped distribution covering six orders of dynamic range of MS signals (Methods, Supplementary Figure 1). The most highly expressed proteins include the core histones, tubulins as well as heat shock proteins. Both the most abundant proteins detected as well as the lowest ones have been previously reported in other studies with a similar approach (43). We kept for further analysis only those proteins containing at least two valid quantification values over the 3 replicates in each cell line. This very conservative approach avoids imputing missing values, and ensures the results of the statistical tests. Doing so, we used for subsequent analyses the quantitative expression data of 1229 proteins (Supplementary Table 1).

A similar strategy was used to identify and quantify phosphopeptides (Methods). We identified 3746 phosphosites, of which 563 were kept for expression analysis considering the strong filters we defined (Supplementary Table 2). These 563 phosphosites correspond to 381 proteins. Overall, 135 proteins were associated with quantitative expression values both at the proteomic and phosphoproteomic levels, with a correlation ranging from 0.43 to 0.62 in each of the four cell lines (Supplementary Figure 2). Therefore, the level of phosphorylation of a protein is not strictly correlated to its level of expression, but might also reflect its activity status.

The unsupervised clustering of the quantified proteins and phosphosites first confirms that the cell line replicates cluster together (Figure 1, D and E). In addition, we observed that the benign PNT1A cell line clusters with the resistant DU145. The genetic instability associated with continuous propagation in culture is a particular problem with benign immortalized cell lines such as PNT1A, in which the insertion of viral DNA drives the cell to replicate continuously (44). This might explain why its global expression patterns may be similar to that of more malignant cell lines.

### The highly-expressed housekeeping proteome

A large number of proteins are essential in all the cells, suggesting that their expression is crucial for the maintenance of basic functionality and survival (45). These proteins are often called housekeeping. We focused here on the top 10% most expressed proteins in each cell line, corresponding to 321 proteins. Among those 321 highly expressed proteins, 257 are common to the four cell lines (Methods, Supplementary Table 3). This means that 80% of the most expressed proteins are the same in all the four cell lines studied here, and can thereby be defined as the highly-expressed housekeeping proteome.

This housekeeping proteome is enriched in functions related to RNA metabolism and response to oxidative stress (functional enrichments with G:Profiler (29), Methods and Supplementary Table 5). It contains for instance many RNA binding proteins (mainly from the RPS family) and structural constituents of the ribosome. Eight members of the eukaryotic chaperonin TriC/CCT complex are also highly abundant in all the four cell lines studied.

### LNCaP, DU145 and PC3 cancer cell lines characterization

In a second step, we focused on the differences between the cell lines. We first conducted an ANOVA analysis to identify the proteins and phosphosites with the most variation among the four cell lines (Methods). 46 proteins and 13 phosphosites (corresponding to 13 proteins) are varying significantly among the four cell lines (FDR < 0.01, Supplementary Tables 3 and 4). Almost half of the 46 ANOVA-significant proteins play a role in stress response (e.g., DNAJB1, VDAC1, ZYX, TCEB1), several are involved in actin cytoskeleton organization (e.g., ACTN1, RHOA, PLS3), and 15 proteins are associated with RNA binding (e.g., CCT6A, NOP2, OCT3, HNRNPA2B1). Among the 13 proteins with phosphosites associated with ANOVA-significant variations in the four cell lines, five are cell-adhesion molecule binding (SEPT9, AHNAK, TNKS1BP1, SCRIB, TAGLN2). Of note, Septin-9 (SEPT9), a filament-forming cytoskeletal GTPase, presents significant variations across the cell lines both at the protein and Serine-30 phosphosite levels (Supplementary Figure 3). SEPT9 has been shown to be highly expressed in PC and positively correlates with malignant progression (46).

Interestingly, two highly expressed housekeeping proteins are associated with phosphosites differentially expressed between the four cell lines according to the ANOVA analysis. First, TAGLN2 presents a significant variation in the Serine163 expression. In liver cancer, this protein has been reported as a putative tumor suppressor and the involvement of its phosphorylation in actin binding and cell migration has been demonstrated (47). Second, HNRNPA1, involved in the packaging of pre-mRNA, is highly expressed in the four cell lines, but also shows significant differential phosphorylation levels in the Serine-6. To our knowledge, a role for HNRNPA1 phosphorylation in PC has not been described previously.

In order to provide insights into the cellular mechanisms that are involved in cell malignant transformation, we then compared protein and phosphosite levels in each of the three cancer cell lines (LNCaP, DU145 and PC3) to the benign PNT1A cell line (Methods). On a global scale, LNCaP clusters apart and appears to be the most divergent cell line (Figure 1, D). LNCaP cells display 226 up- and 219 downregulated proteins as compared to PNT1A (Supplementary Table 3). Functional enrichment analyses reveal that the proteins upregulated in LNCaP are related to cellular metabolism (Figure 2, A, Supplementary Table 5). The association of tumorigenesis and metabolism is well established; it is not surprising that a cancer cell, in order to meet its increased requirements of proliferation, displays fundamental changes in pathways of energy metabolism and nutrient uptake (48). In contrast, the proteins downregulated in LNCaP as compared to PNT1A are enriched in cell recognition and protein/RNA localization processes. Protein and RNA localization mechanisms have shown to play pivotal roles for the presence of specific protein components in cancer cell protrusions, involved in cell migration and invasion (49). Cell recognition is one of the ways that cells communicate with each other and their environment (adhesion proteins, surface molecules); loss of cell recognition has been shown to lead to cancer development (50). IPA analysis (Methods) confirmed a high metabolic activity in LNCaP, in particular an upregulation of TCA cycle for aerobic respiration. It further delineates a downregulation in the RAN signaling pathway, central to the nucleocytoplasmic transport, with seven downregulated proteins, including RAN and its regulator RANBP1, four importins and one exportin (Supplementary Table 5).

**Fig. 2.**
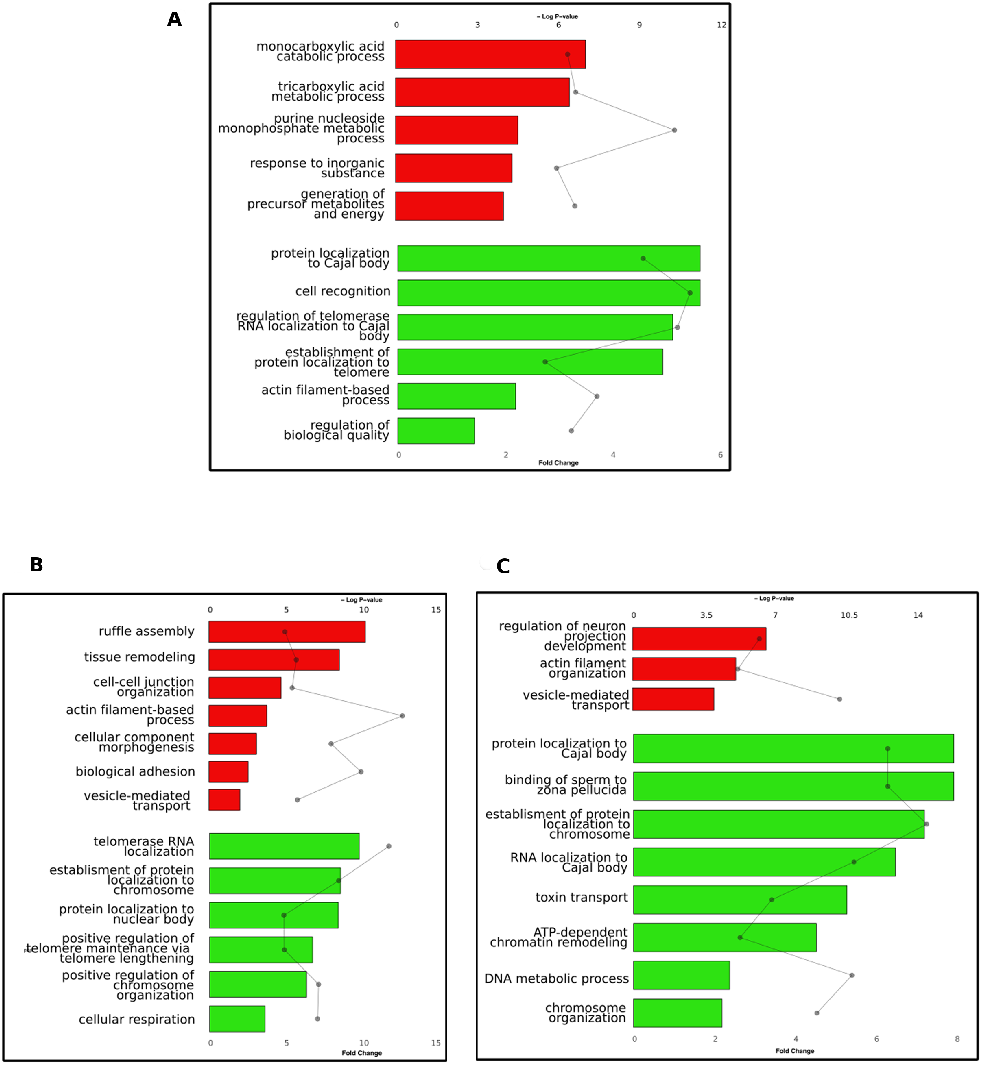
Functional Enrichments of proteins up- and downregulated in PC cell lines. Bar graphs represent relative fold change of Gene Ontology Biological Processes among (A) LNCaP, (B) DU145, (C) PC3 upregulated proteins (red bars) and downregulated proteins (green bars), as compared to PNT1A cells. Significance is represented in the dot plot by −log (P-values).

The resistant cell line DU145 presents 80 up- and 92 downregulated proteins as compared to PNT1A. Upregulated proteins are enriched in transport and cellular organization processes. Moreover, 61/80 proteins upregulated in DU145 are annotated as extracellular proteins. By contrast, we observed that proteins downregulated in DU145 as compared to PNT1A are enriched in cellular respiration and protein/RNA localization (Figure 2, B, Supplementary Table 5). IPA analysis confirmed an upregulation of actin and Rho signaling and a downregulation of TCA cycle for aerobic respiration.

Finally, the most tumorigenic cell line, PC3, displays 180 up- and 158 downregulated proteins as compared to PNT1A. The upregulated proteins are enriched in vesicle-mediated transport, as it is the case for the other resistant cell line DU145 (Figure 2, C, Supplementary Table 5). In recent years, several publications have proposed vesicle-mediated transport as a mechanism to explain the transfer of resistance to drugs among tumorigenic cells (51). In addition, many proteins upregulated in PC3 are localized in the extracellular exosome. The proteins downregulated in PC3 are enriched in toxin transport and protein-RNA localization processes. These functional enrichments are complemented by the IPA analysis that revealed strong enrichment in epithelial adherence junction annotation among the upregulated proteins in PC3. Overall, we identified 13 proteins upregulated and 19 proteins downregulated together in LNCaP, DU145 and PC3 cells as compared to PNT1A (Supplementary Table 3). We propose that these proteins, differentially expressed in the PC cell lines as compared to the benign cell line, could constitute markers of oncogenic transformation. The upregulated proteins are almost all annotated for secretion and exosomes (e.g., RAB5B, RAB7A, RPL36A, NES, SRI). It has been recently described that exosomes derived from PC cells modulate the prostatic tumor adjacent environment by inducing, among others, tumor-associated target cells growth (52). Among the 19 downregulated proteins, several are annotated for regulation of protein stability and chaperone-mediated protein folding, and almost half are involved in DNA metabolism. Overall, many proteins of the chaperonin TriC/CCT folding complex, which were observed as highly abundant in all cell lines and thereby classified as housekeeping, are also underexpressed in the three cancer cell lines as compared to PNT1A. The TriC/CCT chaperonin complex directly modulates the folding and activity of as many as 10% of cytosolic client proteins (53, 54). Recently, the TRiC/CCT complex was also shown to be required for maintaining the wild-type conformation of the tumor suppressor p53 (55). The downregulation of this chaperone complex could promote the oncogenic functions of p53, such as cancer cell migration and invasion.

We reproduced the cell line characterization protocol for phosphosites, thereby identifying 146 up- and 98 downregulated phosphosites in LNCaP, 5 up- and 3 down in DU145, and 82 up- and 44 down in PC3, as compared to PNT1A. No functional enrichments were significant for the corresponding proteins. Nevertheless, two proteins are associated with phosphosites significantly deregulated in all three PC cell lines as compared to PNT1A. First, TP53BP1 (tumor protein p53 binding protein 1) phosphosites Serine-500 and Threonine-1056 are downregulated in LNCaP. TP53BP1 Serine-500 phosphosite is also downregulated in DU145, and the Threonine-1056 phosphosite downregulated in PC3, as compared to PNT1A. This TP53BP1 protein is well known to be involved in DNA Damage Response (DDR) and its phosphorylation could be a marker of malignant transformation (56). Previously published studies described TP53BP1 phosphorylation necessary for recruitment to DNA double strand breaks (DSB) (57). In this context, a downregulation of TP53BP1 phosphorylation, as we observed in the three PC cell lines, could lead to impaired DDR. Second, the DEADbox RNA helicase 10 (DDX10) Serine-539 phosphosite is significantly upregulated in LNCaP, DU145 and PC3 as compared to PNT1A. DDX10 is an ATP-dependent RNA helicase (58), but, to our knowledge, little is known about its phosphorylation and function in cancer. Other members of the same family of RNA helicases have been well described, and the phosphorylation of DDX p68 is reported to be associated with cancer development and cell proliferation (59). Interestingly, the phosphosite Serine-539 that we identified as upregulated in PC cell lines is one of the known post-translational DDX modification sites (60). Thus, our approach allowed us identifying a well-known cancer-related phosphosite, as well as another potential new candidate.

### Identification of Resistance markers

One of the features of PC is, in most cases, its progression to highly aggressive and incurable castration resistant (CR) disease after androgen deprivation therapy. Identifying resistance biomarkers is essential to guide the development of new therapeutic strategies and avoid drug resistance. In order to identify proteins and processes potentially involved in resistance, we compared the protein expression levels in LNCaP cell line (castration-sensitive, CS) with DU145 and PC3, the two castration-resistant cell lines (CR). We found 135 proteins upregulated and 135 downregulated in CR as compared to CS cell lines, and propose them as resistance biomarkers (Supplementary Table 3). Protein biomarkers upregulated in the CR contexts are functionally enriched in processes related to cell-cell adhesion and external communication (Figure 3, A, Supplementary Table 5). This finding is in accordance with previously published studies demonstrating the involvement of these processes in invasion and metastasis, features for which CR cells have a higher potential (61). Conversely, proteins downregulated in CR are enriched in cellular respiration and protein maturation processes. The downregulation of cellular respiration in the CR context could highlight the Warburg effect (62), in which castration-resistant progression would be associated with a switch from oxidative respiration to glycolysis as primary energy source. The ROMA pathway analysis tool (30) also points to a downregulation in CR cells of oxidative phosphorylation and metabolic pathways such as fatty acid metabolism, as well as signaling pathways related to p53 and apoptosis (Figure 3, B). Conversely, it reveals an upregulation of the epithelial-mesenchymal transition (EMT) and reactive oxygen species (ROS) pathways. EMT refers to the morphological and functional alterations involved in cancer invasion (63). Finally, IPA analysis points to an upregulation of actin cytoskeleton and Rho signaling in CR cells, and further identifies an upregulation of Integrin Signaling and Calpain protease signaling.

**Fig. 3.**
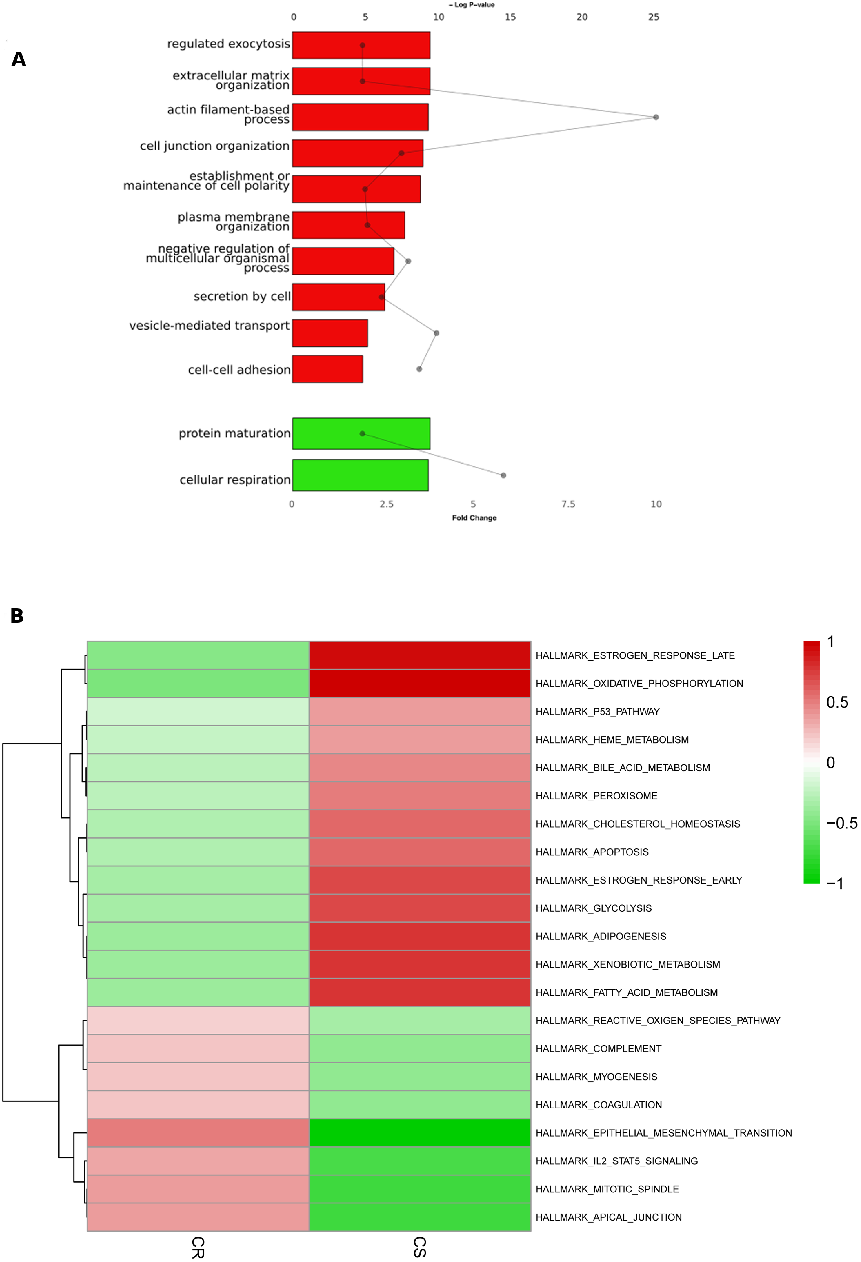
Functional Enrichments of protein resistance biomarkers. (A) Bar graphs represent relative fold change of Gene Ontology Biological Processes among proteins upregulated (red bars) and downregulated (green bars) in Castration Resistant cell lines DU145 and PC3 as compared to castration-sensitive LNCaP cell line. Significance is represented in the dot plot by −log (P-values). (B) Clustered heatmap of ROMA pathway analysis. The color intensities correspond to the values of the scores of each signaling pathway (red, upregulated; green, downregulated).

Phosphoproteomics data reveal 41 phosphosites upregulated and 40 downregulated in CR versus CS, which we also predict as resistance biomarkers (Supplementary Table 4). The 41 upregulated phosphosites concern essentially nuclear proteins involved in functions such as transcription regulation, genome stability and RNA processing (e.g., SMARCC1, SRRM1, SRRM2, SSB). The deregulation of these processes, and their implication in cancer development and progression, have been largely documented (64). Moreover, 2 kinases are hyper-phosphorylated in the resistant context. First, the Serine/threonine-protein kinase N2 (PKN2), which plays a role in the regulation of cell cycle progression, actin cytoskeleton assembly, cell migration, cell adhesion, tumor cell invasion and transcription activation signaling processes. It was recently shown to be phosphorylated by the PI-3 Kinase pathway and implicated in prostate cancer progression (65). Second, the nuclear receptor binding protein (NRBP1), which is involved in subcellular ER-Golgi trafficking. To our knowledge, a role of its phosphorylation status in prostate cancer has not been described previously.

The 40 downregulated phosphosites concern mainly proteins involved in cell migration and invasion, such as PLEC, AHNAK, ESYT1 and ZYX. A group of kinases sharing the same identified peptide and that consequently cannot be distinguished with the MS experiment (CDK2;CDK3;CDK1;CDC2) shows a decrease in phosphorylation activity in the CR context.

Kinase-Substrate Enrichment Analysis (KSEA (33), Methods) predicted the high activity of 3 kinases, namely CDK1, MAPK13 and MAPK3, with 9, 4 and 3 targeted phosphosites that present significant changes in the CR context, respectively (Supplementary Table 4). For instance, the Serine-25 and Serine-38 of the stathmin protein (STMN1) are targets of the three kinases. The STMN1 protein displays a complex pattern of activity and phosphorylation in cancers (66). The sequestosome 1 protein (SQSTM1) Threonine-269 and Serine-272 are targets of both CDK1 and MAPK13.

Another interesting set of putative biomarkers can be derived from the proteins and phosphosites that have been identified in the MS experiment, but that were not further considered for quantification analyses because of the strong filtering criteria we have defined. We thus rescued the proteins and phosphosites that have been identified in at least 2 replicates in the CS cell line but that are completely absents in the CR cell lines, and vice-versa (Methods). This concerns 140 proteins and 5 phosphosites that are identified only in the CR cell lines, and 8 proteins and 108 phosphosites that are identified only in the CS cell line. Focusing particularly on kinases, 8 of them are identified only in the CR cell lines (CALM1, EGFR, EIF2AK2, EPHA2, HK2, PIK3R4, PPP4C, ROCK2). A majority of these kinases are involved in response to stress. Two other kinases are associated with phosphosites identified only in the CR contexts (PRPF4B, TAOK1). TAOK1 is particularly appealing as it activates the Hippo pathway involved in cellular homeostasis (67). Finally, it is to note that some phosphosites associated to significantly different levels of phosphorylation are found in proteins that are quantified by our approach, and not differentially expressed in the ANOVA. These might represent functionally relevant candidates. These include 12 proteins (DKC1, BCLAF1, SRRM2, NAP1L4, CLNS1A, TJP1, API5, SSB, SQSTM1, DHCR7, NCBP1).

### Proteome and phosphoproteome integration in a molecular network

We finally sought to provide a larger-scale functional interpretation of resistance-associated candidate biomarkers. The separated analysis of the proteomics and phosphoproteomics datasets provided one-dimensional views of cellular processes. We expect to obtain a comprehensive perspective of cellular processes and their interplays by integrating the information about protein abundances, activation status and molecular interactions (68, 69). Toward this goal, we devised a network-guided integration of CS and CRPC cell lines proteome and phosphoproteome, by mapping the candidate biomarkers to molecular complex interaction data (Methods). The resulting network is composed of 356 nodes and 1161 edges, including a large connected component encompassing 194 nodes and 1098 edges (Figure 4). The network reveals the links between upand downregulated proteins, upand downregulated phosphosites and corresponding proteins, as well as the links between the proteins and phosphosites that were identified by the MS approach only in the CR or CS contexts. At-a-glance, we can observe that the network is organized around several strongly connected subnetworks.

**Fig. 4.**
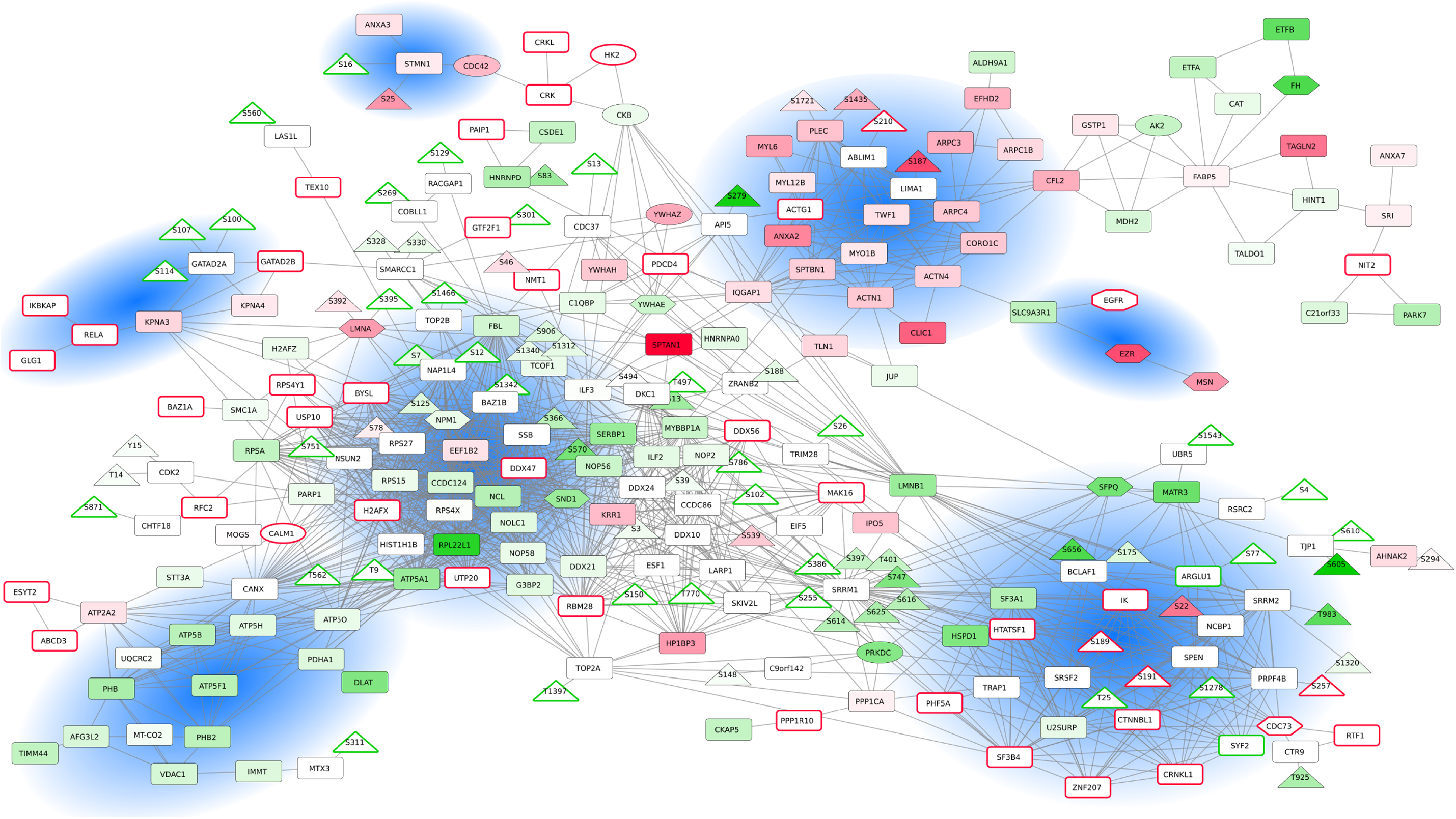
Network of CR biomarker interactions. Proteins (boxes) and phosphosites (triangles) significantly upregulated or downregulated in the CR contexts are mapped in red or green, respectively, with color intensities related to fold-changes. For visualization purposes, the expression values correspond to the mean of the expression of PC3 and DU145 cell lines. Proteins and phosphosites identified only in CR (DU145 and PC3) or CS (LNCaP) cells lines are squared in red and green, respectively.

First, we identified a cell migration/invasion subnetwork, which is composed mainly of upregulated proteins in CR cells (e.g., ANXA2, IQGAP1, ACTN4, TWF1, MYO1B, CORO1C, ARPC4) (Figure 4). It contains in particular the plectrin protein (PLEC), overexpressed and hyperphosphorylated in CR; this protein is known to interlink cytoskeleton elements and promote cancer cell invasion and migration (70). Indeed, it was shown that along with vimentin intermediate filaments, plectrin provide a scaffold for invadopodia formation, facilitating cancer cell invasion extravasation for metastasis (71). Recently, (72) demonstrated that upregulation of vimentin and plectrin expressions positively correlates with the invasion and metastasis of androgen-independent PC cells. Another interesting member of this complex is ACTG1 (actin gamma-1), which is not identified, and thereby might be not expressed, in CS cells. ACTG1 is involved in cell motility/cytoskeleton maintenance and cancer cell migration. ACTG1 was shown to induce cancer cell migration in lung cancer cells and hepatocellular carcinoma cells (73). To date, there is no report concerning ACTG1 involvement in PC. The subnetwork also contains components of the Arp2/3 complex (ARPC1B, ARPC3, ARPC4) involved in the regulation of actin polymerization.

A smaller subnetwork, composed of interactions between EZR, MSN, SLC9A3R1 and EGFR, is located close to the larger migration subnetwork. Ezrin (EZR) and moesin (MSN) are scaffolding proteins that are involved in crosslinking cytoskeletal and membrane proteins. Ezrin is involved in oncogenesis through these interactions (74), and it was also shown recently that Ezrin can increase the oncogenic functions of EGFR (75). SLC9A3R1 is also a scaffold protein that connects plasma membrane proteins with members of the ezrin/moesin/radixin family linking them to the actin cytoskeleton and regulating their surface expression (76).

We also identified a small subnetwork of interacting proteins involved in actin cytoskeleton regulation (e.g., STMN1, CDC42, CRLK1). Intriguingly, we found that stathmin1 (STMN1) was both hyper- and hypo-phosphorylated in CR cells. This protein is associated with cancer metastasis and exhibits a complicated phosphorylation pattern in response to various extracellular signals (77).

We next focused on a small subnetwork composed of proteins underexpressed in the CR context. It contains prohibitin (PHB), a putative tumor suppressor protein involved in the inhibition of DNA synthesis and regulating proliferation, and prohibitin-2 (PHB2), a mediator of transcriptional repression by nuclear receptors, also potentially involved in mitochondrial respiration. Indeed, the subnetwork also contains the VDAC1 mitochondrial membrane and plasma membrane channel, involved in apoptosis. The role of this subnetwork is unclear, but the proteins are depicted as members of the same complexes in the Hu.map dataset (35). The subnetwork is tightly linked to another subnetwork containing many mitochondrial membrane ATP synthase proteins (e.g., ATP5F1, ATP5B, ATP5H), also downregulated in CR cell lines.

A heterogeneous subnetwork is composed of many proteins involved in splicing and RNA processing, that are either up- or downregulated in CR cells (Figure 4). Splicing events control gene expression and their alterations have been shown to play a role in cancer (78) and specifically in PC (79). Fine regulation of expression and/or phosphorylation status determines whether a splicing factor functions as a splicing repressor or activator (80, 81). The subnetwork contains the hypo-phosphorylated splicing factors SRRM1 (a highly phosphorylated protein under normal conditions (82)) and SRRM2. It also contains NCBP1, which is identified as hyper-phosphorylated, and PRPF4B kinase and SPEN that were both hypo-phosphorylated. The subnetwork also incorporates pre-mRNA splicing factor SYF2, absent in CR cells, and SF3A1, TRAP1 and HSPD1 proteins that are downregulated in CR cells. The protein phosphatase 1 (PPP1CA) is contrarily upregulated. Interestingly, we can also observe many proteins identified in the CR cell lines and absent in the CS cell line, all involved in RNA processing and splicing (IK, ZNF207, CTNNBL1, CRNKL1, SF3B4, HTATSF1, PHF5A, PPP1R10). PPP1R10, the Ser/Thr-protein phosphatase-1 regulatory subunit 10 is only expressed in CR cells and is absent in CS cells. It has been shown that certain Ser/Thr-specific protein phosphatases are required for catalytic steps of pre-mRNA splicing (83).

We then emphasize a large and highly connected component (Figure 4) composed of proteins implicated in DNA damage response. It contains protein biomarkers downregulated in CR cells (NPM1, NOLC1, RPL22L1, FBL, G3BP2), but also several proteins identified only in the CR cell lines, namely H2AFX, kinase CALM1, DDX47, UTP20, USP10, BYSL. All these proteins interact with single-strand DNA-binding protein and are involved in DNA repair and genome stability (84). DNA repair and DNA damage response are known to be defective in PC and lead to genome instability (85). Interestingly, several of the proteins of this subnetwork (e.g., UTP20, BYSL, RPL22L1, NOLC1) are known for their role in RNA processing. There is an increasing number of studies demonstrating the involvement of RNA processing factors in DNA damage response (86, 87). For instance, NOLC1 (nuclear and coiled-body phosphoprotein-1) is a regulator of RNA polymerase I and has been recently shown to regulate the nucleolar retention of TERF2, inducing telomeric DNA damage (88).

A closer look into this molecular network allowed us to pinpoint several interesting smaller subnetworks. For instance, we noticed a small subnetwork composed of interacting proteins RELA, IKBKAP, GLG1, KPNA3, KPNA4. Importin subunits alpha-4 (KPNA3) and alpha-3 (KPNA4) are involved in nuclear transport of NF kappa B (89), and an elevated activity of the NF-kappa B signaling in CRPC is positively correlated with poor prognosis in CRPC (90). Close to this subnetwork, GATAD2B is known to form a homodimer with GATAD2A and the complex is part of a highly conserved chromatin-remodeling complex, the NuRD complex associated with DNA damage-induced transcription repression but also metastasis and EMT (91, 92). This subnetwork is also linked to the SWI/SNF complex subunit SMARCC1, which contains downregulated phosphosites in PC3 and DU145 cells. SMARCC1 positively regulates transcription and was previously shown to induce PC survival (93). It interacts with proteins associated with phosphosites only detected in CS cells (transcriptional elongation factor TRIM28, transcription kinase BAZ1B and TOP2B), as well as with proteins only identified in PC3 and DU145 cells (e.g., GATAD2B, GTF2F1), all involved directly or indirectly in transcription regulation. The transcriptional reprogramming in PC progression has been extensively studied, as it is one of the hallmarks of CRPC (3, 94–96).

## Discussion

We here generated and explored a SILAC proteomics and phosphoproteomics dataset of prostate cell lines. We selected the PNT1A, LNCaP, DU145 and PC3 cell lines first because they are frequently used in prostate cancer research (97, 98), and molecular profiling and comparisons would thereby be highly valuable for researchers using them in routine. In addition, the elected cell lines are representative of normal, cancerous and castration-resistant prostate tissue, and therefore reflect progression of the disease.

We decided to monitor proteome and phosphoproteome jointly as they can give a complementary picture of the molecular dynamics of the cells. Phosphoproteome characterization provides insights into proteins’ phosphorylation levels, which are not strictly correlated to proteins level of expression, but also reflect protein activity status (99). Molecular characterization at these two levels of information is a clear advantage in our study. We identified 3 219 proteins at 1% FDR, and after several filtering steps, we performed subsequent functional explorations on 1229 proteins. We elected this conservative approach in order to avoid imputation of missing values and ensure the results of the statistical analyses. On the phosphoproteomics side, we identified 3746 phosphosites among which 563 are used for subsequent analyses. We explored the proteins and phosphosites associated to the four PC cell lines; these profiles and associated cellular processes could be related to underlying biology of the cell lines (Figure 1, Supplementary Tables 1-3, Supplementary Figure 1). We further compared protein and phosphosite levels in each of the three cancer cell lines (LNCaP, DU145 and PC3) to the benign PNT1A cell line (Figure 2, Supplementary Tables 3-5). Doing so, we proposed several candidates that could constitute markers of oncogenic transformation. Notably, two proteins are associated with phosphosites significantly deregulated in all three PC cell lines as compared to PNT1A: TP53BP1, well-known cancer-related phosphosites downregulated in all three PC cell lines and DDX10, a potential new candidate, upregulated in PCa cell lines. We also highlighted differentially expressed proteins and processes potentially involved in resistance, by comparing the sensitive LNCaP cell line to the two resistant DU145 and PC3 cell lines (Figure 3). Interestingly, we identified 12 proteins associated to significant differential phosphorylation but not different protein levels, including several RNA binding proteins, sequestosome-1 protein (involved in autophagy in relation to many crucial signalling pathways), as well as DHCR7, an enzyme involved in cholesterol metabolism, and TJP1, a tight junction protein. Finally, we proposed an integrated mapping of protein abundances, activation status and molecular interactions, towards functional interpretation of resistance-associated candidate biomarkers (Figure 4). We observed that the network is organized around several strongly connected subnetworks (cell migration/invasion subnetwork, actin cytoskeleton regulation, DNA synthesis and regulating proliferation, splicing and RNA processing, DNA damage response) and several interesting smaller subnetworks (transcription regulation, nuclear transport). Several of these proteins and phosphosites are already related to cancer resistance in general, and some specifically to PCa. Overall, this analysis represents a valuable resource that could be used as a starting point for further hypothesis and experimental investigations.

## Conclusion

The complex nature of PC, due to its clinical and molecular heterogeneities, makes it difficult to determine a perfect model representing tumor development, and precludes easy correlation of carcinoma cell lines with specific stages of PC. Nevertheless, PC cell lines routinely used for the last three decades have provided valuable resources for understanding important functional molecular mechanisms involved in this disease. In the present study, we used four cell lines that constitute a gold standard for pre-clinical studies of PC progression. We conducted a large SILAC-based Mass Spectrometry identification and quantification of peptides and phosphopeptides of prostate benign, castration-sensitive (CS) and castration-resistant (CR) cells, and characterized housekeeping, cell line, cancer and resistance associated proteomes and phosphoproteomes.

## ACKNOWLEDGEMENTS

This work was supported by the French “Plan Cancer 2009-2013” (Systems Biology call). The authors thank Christine Brun, Andreas Zanzoni and all the partners of the Hsp27BioSys project for fruitful discussion. The differential proteomic analyses were done using the Mass Spectrometry facility of Marseille Proteomics (http://map.univmed.fr/) supported by IBISA (Infrastructures Biologie Santé et Agronomie), the Cancéropôle PACA, the Provence-Alpes-Côte d’Azur Region, the Institut Paoli-Calmettes and the Centre de Recherche en Cancérologie de Marseille.

## Notes

#### Summary of Updates

Sentence added in the conclusion of the abstract to emphasise some of the article results. Discussion section added to the article.

## Bibliography

1. A. Fusi, G. Procopio, S. Della Torre, R. Ricotta, G. Bianchini, R. Salvioni, L. Ferrari, A. Martinetti, G. Savelli, S. Villa, and E. Bajetta. Treatment options in hormone-refractory metastatic prostate carcinoma. Tumori, 90(6):535–46, 2004. Fusi, Alberto Procopio, Giuseppe Della Torre, Silvia Ricotta, Riccardo Bianchini, Gianpaolo Salvioni, Roberto Ferrari, Leonardo Martinetti, Antonia Savelli, Giordano Villa, Sergio Bajetta, Emilio Review Italy Tumori Tumori. 2004 Nov-Dec;90(6):535–46.

2. T. Karantanos, C. P. Evans, B. Tombal, T. C. Thompson, R. Montironi, and W. B. Isaacs. Understanding the mechanisms of androgen deprivation resistance in prostate cancer at the molecular level. Eur Urol, 67(3):470–9, 2015. Karantanos, Theodoros Evans, Christopher P Tombal, Bertrand Thompson, Timothy C Montironi, Rodolfo Isaacs, William B P30 CA016672/CA/NCI NIH HHS/United States Review Switzerland European urology Eur Urol. 2015 Mar;67(3):470–9. doi: 10.1016/j.eururo.2014.09.049. Epub 2014 Oct 8.

3. M. Katsogiannou, H. Ziouziou, S. Karaki, C. Andrieu, M. Henry de Villeneuve, and P. Rocchi. The hallmarks of castration-resistant prostate cancers. Cancer Treat Rev, 41(7):588–97, 2015. Katsogiannou, Maria Ziouziou, Hajer Karaki, Sara Andrieu, Claudia Henry de Villeneuve, Marie Rocchi, Palma Research Support, Non-U.S. Gov’t Review Netherlands Cancer treatment reviews Cancer Treat Rev. 2015 Jul;41(7):588–97. doi: 10.1016/j.ctrv.2015.05.003. Epub 2015 May 9.

4. M. D. Galsky, A. C. Small, C. K. Tsao, and W. K. Oh. Clinical development of novel therapeutics for castration-resistant prostate cancer: historic challenges and recent successes. CA Cancer J Clin, 62(5):299–308, 2012. Galsky, Matthew D Small, Alexander C Tsao, Che-kai Oh, William K Review United States CA: a cancer journal for clinicians CA Cancer J Clin. 2012 Sep-Oct;62(5):299–308. doi: 10.3322/caac.21141. Epub 2012 Apr 24.

5. D. L. Suzman and E. S. Antonarakis. Castration-resistant prostate cancer: latest evidence and therapeutic implications. Ther Adv Med Oncol, 6(4):167–79, 2014. Suzman, Daniel L Antonarakis, Emmanuel S Review England Therapeutic advances in medical oncology Ther Adv Med Oncol. 2014 Jul;6(4):167–79. doi: 10.1177/1758834014529176.

6. D. Iglesias-Gato, E. Thysell, S. Tyanova, S. Crnalic, A. Santos, T. S. Lima, T. Geiger, J. Cox, A. Widmark, A. Bergh, M. Mann, A. Flores-Morales, and P. Wikstrom. The proteome of prostate cancer bone metastasis reveals heterogeneity with prognostic implications. Clin Cancer Res, 24(21):5433–5444, 2018. Iglesias-Gato, Diego Thysell, Elin Tyanova, Stefka Crnalic, Sead Santos, Alberto Lima, Thiago S Geiger, Tamar Cox, Jurgen Widmark, Anders Bergh, Anders Mann, Matthias Flores-Morales, Amilcar Wikstrom, Pernilla United States Clinical cancer research: an official journal of the American Association for Cancer Research Clin Cancer Res. 2018 Nov 1;24(21):5433–5444. doi: 10.1158/1078-0432.CCR18-1229. Epub 2018 Jul 24.

7. D. Iglesias-Gato, P. Wikstrom, S. Tyanova, C. Lavallee, E. Thysell, J. Carlsson, C. Hagglof, J. Cox, O. Andren, P. Stattin, L. Egevad, A. Widmark, A. Bjartell, C. C. Collins, A. Bergh, T. Geiger, M. Mann, and A. Flores-Morales. The proteome of primary prostate cancer. Eur Urol, 69(5):942–52, 2016. Iglesias-Gato, Diego Wikstrom, Pernilla Tyanova, Stefka Lavallee, Charlotte Thysell, Elin Carlsson, Jessica Hagglof, Christina Cox, Jurgen Andren, Ove Stattin, Par Egevad, Lars Widmark, Anders Bjartell, Anders Collins, Colin C Bergh, Anders Geiger, Tamar Mann, Matthias Flores-Morales, Amilcar Research Support, Non-U.S. Gov’t Switzerland European urology Eur Urol. 2016 May;69(5):942–52. doi: 10.1016/j.eururo.2015.10.053. Epub 2015 Dec 2.

8. H. Kuruma, S. Egawa, M. Oh-Ishi, Y. Kodera, and T. Maeda. Proteome analysis of prostate cancer. Prostate Cancer Prostatic Dis, 8(1):14–21, 2005. Kuruma, H Egawa, S Oh-Ishi, M Kodera, Y Maeda, T Research Support, Non-U.S. Gov’t Review England Prostate cancer and prostatic diseases Prostate Cancer Prostatic Dis. 2005;8(1):14–21.

9. H. Kuruma, S. Egawa, M. Oh-Ishi, Y. Kodera, M. Satoh, W. Chen, H. Okusa, K. Matsumoto, T. Maeda, and S. Baba. High molecular mass proteome of androgen-independent prostate cancer. Proteomics, 5(4):1097–112, 2005. Kuruma, Hidetoshi Egawa, Shin OhIshi, Masamichi Kodera, Yoshio Satoh, Mamoru Chen, Weigiang Okusa, Hiroshi Matsumoto, Kazumasa Maeda, Tadakazu Baba, Shiro Research Support, Non-U.S. Gov’t Germany Proteomics Proteomics. 2005 Mar;5(4):1097–112.

10. D. K. Ornstein and D. R. Tyson. Proteomics for the identification of new prostate cancer biomarkers. Urol Oncol, 24(3):231–6, 2006. Ornstein, David K Tyson, Darren R Review United States Urologic oncology Urol Oncol. 2006 May-Jun;24(3):231–6.

11. L. N. Johnson. The regulation of protein phosphorylation. Biochem Soc Trans, 37(Pt 4):627–41, 2009. Johnson, Louise N G0400957/Medical Research Council/United Kingdom Biotechnology and Biological Sciences Research Council/United Kingdom Medical Research Council/United Kingdom Wellcome Trust/United Kingdom Research Support, Non-U.S. Gov’t Review England Biochemical Society transactions Biochem Soc Trans. 2009 Aug;37(Pt 4):627–41. doi: 10.1042/BST0370627.

12. C. Cans, R. Mangano, D. Barila, G. Neubauer, and G. Superti-Furga. Nuclear tyrosine phosphorylation: the beginning of a map. Biochem Pharmacol, 60(8):1203–15, 2000. Cans, C Mangano, R Barila, D Neubauer, G Superti-Furga, G Research Support, Non-U.S. Gov’t Review England Biochemical pharmacology Biochem Pharmacol. 2000 Oct 15;60(8):1203-15.

13. E. Lopez Villar, L. Madero, A. Lopez-Pascual J, and C. Cho W. Study of phosphorylation events for cancer diagnoses and treatment. Clin Transl Med, 4(1):59, 2015. Lopez Villar, Elena Madero, Luis A Lopez-Pascual, Juan C Cho, William Germany Clinical and translational medicine Clin Transl Med. 2015 Dec;4(1):59. doi: 10.1186/s40169-015-0059-0. Epub 2015 May 24.

14. H. C. Harsha and A. Pandey. Phosphoproteomics in cancer. Mol Oncol, 4(6):482–95, 2010. Harsha, H C Pandey, Akhilesh U54 RR020839/RR/NCRR NIH HHS/United States N01HV-28180/HV/NHLBI NIH HHS/United States U54 RR 020839/RR/NCRR NIH HHS/United States U54 RR020839-08/RR/NCRR NIH HHS/United States N01 HV028180/HV/NHLBI NIH HHS/United States N01HV28180/HL/NHLBI NIH HHS/United States Research Support, N.I.H., Extramural Research Support, Non-U.S. Gov’t Review United States Molecular oncology Mol Oncol. 2010 Dec;4(6):482–95. doi: 10.1016/j.molonc.2010.09.004. Epub 2010 Sep 26.

15. R. M. Lescarbeau and D. L. Kaplan. Quantitative analysis of castration resistant prostate cancer progression through phosphoproteome signaling. BMC Cancer, 14:325, 2014. Lescarbeau, Reynald M Kaplan, David L P41 EB002520-05/EB/NIBIB NIH HHS/United States Research Support, N.I.H., Extramural England BMC cancer BMC Cancer. 2014 May 8;14:325. doi: 10.1186/1471-2407-14-325.

16. N. Jiang, K. Hjorth-Jensen, O. Hekmat, D. Iglesias-Gato, T. Kruse, C. Wang, W. Wei Ke, B. Yan, Y. Niu, J. V. Olsen, and A. Flores-Morales. In vivo quantitative phosphoproteomic profiling identifies novel regulators of castration-resistant prostate cancer growth. Oncogene, 34(21):2764–76, 2015. Jiang, N Hjorth-Jensen, K Hekmat, O Iglesias-Gato, D Kruse, T Wang, C Wei, W Ke, B Yan, B Niu, Y Olsen, J V Flores-Morales, A Research Support, Non-U.S. Gov’t England Oncogene Oncogene. 2015 May 21;34(21):2764–76. doi: 10.1038/onc.2014.206. Epub 2014 Jul 28.

17. X. Wang, P. A. Stewart, Q. Cao, Q. X. Sang, L. W. Chung, M. R. Emmett, and A. G. Marshall. Characterization of the phosphoproteome in androgen-repressed human prostate cancer cells by fourier transform ion cyclotron resonance mass spectrometry. J Proteome Res, 10 (9):3920–8, 2011. Wang, Xu Stewart, Paul A Cao, Qiang Sang, Qing-Xiang Amy Chung, Leland W K Emmett, Mark R Marshall, Alan G P01 CA098912/CA/NCI NIH HHS/United States R01 CA122602/CA/NCI NIH HHS/United States Comparative Study Research Support, U.S. Gov’t, Non-P.H.S. United States Journal of proteome research J Proteome Res. 2011 Sep 2;10(9):3920–8. doi: 10.1021/pr2000144. Epub 2011 Jul 26.

18. J. M. Drake, E. O. Paull, N. A. Graham, J. K. Lee, B. A. Smith, B. Titz, T. Stoyanova, M. Faltermeier, V. Uzunangelov, D. E. Carlin, D. T. Fleming, C. K. Wong, Y. Newton, S. Sudha, A. A. Vashisht, J. Huang, J. A. Wohlschlegel, T. G. Graeber, O. N. Witte, and J. M. Stuart. Phosphoproteome integration reveals patient-specific networks in prostate cancer. Cell, 166(4):1041–54, 2016. Drake, Justin M Paull, Evan O Graham, Nicholas A Lee, John K Smith, Bryan A Titz, Bjoern Stoyanova, Tanya Faltermeier, Claire M Uzunangelov, Vladislav Carlin, Daniel E Fleming, Daniel Teo Wong, Christopher K Newton, Yulia Sudha, Sud Vashisht, Ajay A Huang, Jiaoti Wohlschlegel, James A Graeber, Thomas G Witte, Owen N Stuart, Joshua M R01 CA195505/CA/NCI NIH HHS/United States R25 CA098010/CA/NCI NIH HHS/United States R01 CA181242/CA/NCI NIH HHS/United States U54 HG006097/HG/NHGRI NIH HHS/United States R01 GM109031/GM/NIGMS NIH HHS/United States UL1 TR000124/TR/NCATS NIH HHS/United States P30 CA016042/CA/NCI NIH HHS/United States T32 CA009120/CA/NCI NIH HHS/United States R00 CA184397/CA/NCI NIH HHS/United States U24 CA143858/CA/NCI NIH HHS/United States R01 CA180778/CA/NCI NIH HHS/United States P01 CA168585/CA/NCI NIH HHS/United States R01 CA172603/CA/NCI NIH HHS/United States HHMI/HHMI/United States R01 GM089778/GM/NIGMS NIH HHS/United States Research Support, N.I.H., Extramural Research Support, Non-U.S. Gov’t United States Cell Cell. 2016 Aug 11;166(4):1041–54. doi: 10.1016/j.cell.2016.07.007. Epub 2016 Aug 4.

19. T. Geiger, J. Cox, P. Ostasiewicz, J. R. Wisniewski, and M. Mann. Super-silac mix for quantitative proteomics of human tumor tissue. Nat Methods, 7(5):383–5, 2010. Geiger, Tamar Cox, Juergen Ostasiewicz, Pawel Wisniewski, Jacek R Mann, Matthias Research Support, Non-U.S. Gov’t United States Nature methods Nat Methods. 2010 May;7(5):383-5. doi: 10.1038/nmeth.1446. Epub 2010 Apr 4.

20. T. Geiger, J. R. Wisniewski, J. Cox, S. Zanivan, M. Kruger, Y. Ishihama, and M. Mann. Use of stable isotope labeling by amino acids in cell culture as a spike-in standard in quantitative proteomics. Nat Protoc, 6(2):147–57, 2011. Geiger, Tamar Wisniewski, Jacek R Cox, Juergen Zanivan, Sara Kruger, Marcus Ishihama, Yasushi Mann, Matthias Research Support, Non-U.S. Gov’t England Nature protocols Nat Protoc. 2011 Feb;6(2):147–57. doi: 10.1038/nprot.2010.192.

21. A. Shevchenko, M. Wilm, O. Vorm, O. N. Jensen, A. V. Podtelejnikov, G. Neubauer, P. Mortensen, and M. Mann. A strategy for identifying gel-separated proteins in sequence databases by ms alone. Biochem Soc Trans, 24(3):893–6, 1996. Shevchenko, A Wilm, M Vorm, O Jensen, O N Podtelejnikov, A V Neubauer, G Mortensen, P Mann, M England Biochemical Society transactions Biochem Soc Trans. 1996 Aug;24(3):893–6.

22. E. W. Deutsch, A. Csordas, Z. Sun, A. Jarnuczak, Y. Perez-Riverol, T. Ternent, D. S. Campbell, M. Bernal-Llinares, S. Okuda, S. Kawano, R. L. Moritz, J. J. Carver, M. Wang, Y. Ishihama, N. Bandeira, H. Hermjakob, and J. A. Vizcaino. The proteomexchange consortium in 2017: supporting the cultural change in proteomics public data deposition. Nucleic Acids Res, 45(D1):D1100–D1106, 2017. Deutsch, Eric W Csordas, Attila Sun, Zhi Jarnuczak, Andrew Perez-Riverol, Yasset Ternent, Tobias Campbell, David S Bernal-Llinares, Manuel Okuda, Shujiro Kawano, Shin Moritz, Robert L Carver, Jeremy J Wang, Mingxun Ishihama, Yasushi Bandeira, Nuno Hermjakob, Henning Vizcaino, Juan Antonio U54 EB020406/EB/NIBIB NIH HHS/United States P41 GM103484/GM/NIGMS NIH HHS/United States R01 GM087221/GM/NIGMS NIH HHS/United States P50 GM076547/GM/NIGMS NIH HHS/United States Wellcome Trust/United Kingdom England Nucleic acids research Nucleic Acids Res. 2017 Jan 4;45(D1):D1100–D1106. doi: 10.1093/nar/gkw936. Epub 2016 Oct 18.

23. J. Cox, I. Matic, M. Hilger, N. Nagaraj, M. Selbach, J. V. Olsen, and M. Mann. A practical guide to the maxquant computational platform for silac-based quantitative proteomics. Nat Protoc, 4(5):698–705, 2009. Cox, Jurgen Matic, Ivan Hilger, Maximiliane Nagaraj, Nagarjuna Selbach, Matthias Olsen, Jesper V Mann, Matthias England Nature protocols Nat Protoc. 2009;4(5):698–705. doi: 10.1038/nprot.2009.36.

24. J. Cox, N. Neuhauser, A. Michalski, R. A. Scheltema, J. V. Olsen, and M. Mann. Andromeda: a peptide search engine integrated into the maxquant environment. J Proteome Res, 10(4): 1794–805, 2011. Cox, Jurgen Neuhauser, Nadin Michalski, Annette Scheltema, Richard A Olsen, Jesper V Mann, Matthias Research Support, Non-U.S. Gov’t United States Journal of proteome research J Proteome Res. 2011 Apr 1;10(4):1794–805. doi: 10.1021/pr101065j. Epub 2011 Feb 22.

25. J. V. Olsen, S. E. Ong, and M. Mann. Trypsin cleaves exclusively c-terminal to arginine and lysine residues. Mol Cell Proteomics, 3(6):608–14, 2004. Olsen, Jesper V Ong, ShaoEn Mann, Matthias Research Support, Non-U.S. Gov’t United States Molecular & cellular proteomics: MCP Mol Cell Proteomics. 2004 Jun;3(6):608–14. doi: 10.1074/mcp.T400003MCP200. Epub 2004 Mar 19.

26. J. Cox and M. Mann. 1d and 2d annotation enrichment: a statistical method integrating quantitative proteomics with complementary high-throughput data. BMC Bioinformatics, 13 Suppl 16:S12, 2012. Cox, Juergen Mann, Matthias England BMC bioinformatics BMC Bioinformatics. 2012;13 Suppl 16:S12. doi: 10.1186/1471-2105-13-S16-S12. Epub 2012 Nov 5.

27. RCoreTeam. R: A language and environment for statistical computing., 2015.

28. Y. Benjamini and Y Hochberg. Controlling the false discovery rate: A practical and powerful approach to multiple testing. Journal of the Royal Statistical Society Series B (Methodological), Vol. 57(N°1):289–300, 1995.

29. J. Reimand, T. Arak, P. Adler, L. Kolberg, S. Reisberg, H. Peterson, and J. Vilo. g:profiler-a web server for functional interpretation of gene lists (2016 update). Nucleic Acids Res, 44 (W1):W83–9, 2016. Reimand, Juri Arak, Tambet Adler, Priit Kolberg, Liis Reisberg, Sulev Peterson, Hedi Vilo, Jaak England Nucleic acids research Nucleic Acids Res. 2016 Jul 8;44(W1):W83–9. doi: 10.1093/nar/gkw199. Epub 2016 Apr 20.

30. L. Martignetti, L. Calzone, E. Bonnet, E. Barillot, and A. Zinovyev. Roma: Representation and quantification of module activity from target expression data. Front Genet, 7:18, 2016. Martignetti, Loredana Calzone, Laurence Bonnet, Eric Barillot, Emmanuel Zinovyev, Andrei Switzerland Frontiers in genetics Front Genet. 2016 Feb 19;7:18. doi: 10.3389/fgene.2016.00018. eCollection 2016.

31. A. Fabregat, K. Sidiropoulos, P. Garapati, M. Gillespie, K. Hausmann, R. Haw, B. Jassal, S. Jupe, F. Korninger, S. McKay, L. Matthews, B. May, M. Milacic, K. Rothfels, V. Shamovsky, M. Webber, J. Weiser, M. Williams, G. Wu, L. Stein, H. Hermjakob, and P. D’Eustachio. The reactome pathway knowledgebase. Nucleic Acids Res, 44(D1):D481–7, 2016. Fabregat, Antonio Sidiropoulos, Konstantinos Garapati, Phani Gillespie, Marc Hausmann, Kerstin Haw, Robin Jassal, Bijay Jupe, Steven Korninger, Florian McKay, Sheldon Matthews, Lisa May, Bruce Milacic, Marija Rothfels, Karen Shamovsky, Veronica Webber, Marissa Weiser, Joel Williams, Mark Wu, Guanming Stein, Lincoln Hermjakob, Henning D’Eustachio, Peter P41 HG003751/HG/NHGRI NIH HHS/United States U41 HG003751/HG/NHGRI NIH HHS/United States U54 GM114833/GM/NIGMS NIH HHS/United States Research Support, N.I.H., Extramural England Nucleic acids research Nucleic Acids Res. 2016 Jan 4;44(D1):D481–7. doi: 10.1093/nar/gkv1351. Epub 2015 Dec 9.

32. A. Liberzon, C. Birger, H. Thorvaldsdottir, M. Ghandi, J. P. Mesirov, and P. Tamayo. The molecular signatures database (msigdb) hallmark gene set collection. Cell Syst, 1 (6):417–425, 2015. Liberzon, Arthur Birger, Chet Thorvaldsdottir, Helga Ghandi, Mahmoud Mesirov, Jill P Tamayo, Pablo R01 CA121941/CA/NCI NIH HHS/United States R01 CA154480/CA/NCI NIH HHS/United States R01 GM074024/GM/NIGMS NIH HHS/United States U54 CA112962/CA/NCI NIH HHS/United States United States Cell systems Cell Syst. 2015 Dec 23;1(6):417–425. doi: 10.1016/j.cels.2015.12.004.

33. P. Casado, J. C. Rodriguez-Prados, S. C. Cosulich, S. Guichard, B. Vanhaesebroeck, S. Joel, and P. R. Cutillas. Kinase-substrate enrichment analysis provides insights into the heterogeneity of signaling pathway activation in leukemia cells. Sci Signal, 6(268): rs6, 2013 Casado, Pedro Rodriguez-Prados, Juan-Carlos Cosulich, Sabina C Guichard, Sylvie Vanhaesebroeck, Bart Joel, Simon Cutillas, Pedro R BB/G015023/1/Biotechnology and Biological Sciences Research Council/United Kingdom BB/G0115023/1/Biotechnology and Biological Sciences Research Council/United Kingdom G0800914/Medical Research Council/United Kingdom Research Support, Non-U.S. Gov’t United States Science signaling Sci Signal. 2013 Mar 26;6(268):rs6. doi: 10.1126/scisignal.2003573.

34. A. Ruepp, B. Waegele, M. Lechner, B. Brauner, I. Dunger-Kaltenbach, G. Fobo, G. Frishman, C. Montrone, and H. W. Mewes. Corum: the comprehensive resource of mammalian protein complexes–2009. Nucleic Acids Res, 38(Database issue):D497–501, 2010. Ruepp, Andreas Waegele, Brigitte Lechner, Martin Brauner, Barbara Dunger-Kaltenbach, Irmtraud Fobo, Gisela Frishman, Goar Montrone, Corinna Mewes, H-Werner Research Support, Non-U.S. Gov’t England Nucleic acids research Nucleic Acids Res. 2010 Jan;38(Database issue):D497–501. doi: 10.1093/nar/gkp914. Epub 2009 Nov 1.

35. K. Drew, C. Lee, R. L. Huizar, F. Tu, B. Borgeson, C. D. McWhite, Y. Ma, J. B. Wallingford, and E. M. Marcotte. Integration of over 9,000 mass spectrometry experiments builds a global map of human protein complexes. Mol Syst Biol, 13(6):932, 2017. Drew, Kevin Lee, Chanjae Huizar, Ryan L Tu, Fan Borgeson, Blake McWhite, Claire D Ma, Yun Wallingford, John B Marcotte, Edward M DP1 GM106408/GM/NIGMS NIH HHS/United States R01 HD085901/HD/NICHD NIH HHS/United States R21 GM119021/GM/NIGMS NIH HHS/United States R01 HL117164/HL/NHLBI NIH HHS/United States R01 DK110520/DK/NIDDK NIH HHS/United States R35 GM122480/GM/NIGMS NIH HHS/United States F32 GM112495/GM/NIGMS NIH HHS/United States England Molecular systems biology Mol Syst Biol. 2017 Jun 8;13(6):932.

36. P. Shannon, A. Markiel, O. Ozier, N. S. Baliga, J. T. Wang, D. Ramage, N. Amin, B. Schwikowski, and T. Ideker. Cytoscape: a software environment for integrated models of biomolecular interaction networks. Genome Res, 13(11):2498–504, 2003. Shannon, Paul Markiel, Andrew Ozier, Owen Baliga, Nitin S Wang, Jonathan T Ramage, Daniel Amin, Nada Schwikowski, Benno Ideker, Trey P20 GM64361/GM/NIGMS NIH HHS/United States Research Support, Non-U.S. Gov’t Research Support, U.S. Gov’t, Non-P.H.S. Research Support, U.S. Gov’t, P.H.S. United States Genome research Genome Res. 2003 Nov;13(11):2498–504.

37. C. Avances, V. Georget, B. Terouanne, F. Orio, O. Cussenot, N. Mottet, P. Costa, and C. Sultan. Human prostatic cell line pnt1a, a useful tool for studying androgen receptor transcriptional activity and its differential subnuclear localization in the presence of androgens and antiandrogens. Mol Cell Endocrinol, 184(1-2):13–24, 2001. Avances, C Georget, V Terouanne, B Orio, F Cussenot, O Mottet, N Costa, P Sultan, C Comparative Study Research Support, Non-U.S. Gov’t Ireland Molecular and cellular endocrinology Mol Cell Endocrinol. 2001 Nov 26;184(1-2):13–24.

38. P. J. Russell and E. A. Kingsley. Human prostate cancer cell lines. Methods Mol Med, 81: 21–39, 2003. Russell, Pamela J Kingsley, Elizabeth A Review United States Methods in molecular medicine Methods Mol Med. 2003;81:21–39.

39. M. M. Webber, D. Bello, and S. Quader. Immortalized and tumorigenic adult human prostatic epithelial cell lines: characteristics and applications part 2. tumorigenic cell lines. Prostate, 30(1):58–64, 1997. Webber, M M Bello, D Quader, S Research Support, Non-U.S. Gov’t Review United States The Prostate Prostate. 1997 Jan 1;30(1):58–64.

40. S. Mitchell, P. Abel, M. Ware, G. Stamp, and E. Lalani. Phenotypic and genotypic characterization of commonly used human prostatic cell lines. BJU Int, 85(7):932–44, 2000. Mitchell, S Abel, P Ware, M Stamp, G Lalani, E Research Support, Non-U.S. Gov’t England BJU international BJU Int. 2000 May;85(7):932–44.

41. M. Mann. Functional and quantitative proteomics using silac. Nat Rev Mol Cell Biol, 7(12): 952–8, 2006. Mann, Matthias Research Support, Non-U.S. Gov’t England Nature reviews. Molecular cell biology Nat Rev Mol Cell Biol. 2006 Dec;7(12):952–8.

42. S. E. Ong, B. Blagoev, I. Kratchmarova, D. B. Kristensen, H. Steen, A. Pandey, and M. Mann. Stable isotope labeling by amino acids in cell culture, silac, as a simple and accurate approach to expression proteomics. Mol Cell Proteomics, 1(5):376–86, 2002. Ong, Shao-En Blagoev, Blagoy Kratchmarova, Irina Kristensen, Dan Bach Steen, Hanno Pandey, Akhilesh Mann, Matthias Research Support, Non-U.S. Gov’t United States Molecular & cellular proteomics: MCP Mol Cell Proteomics. 2002 May;1(5):376–86.

43. N. Nagaraj, J. R. Wisniewski, T. Geiger, J. Cox, M. Kircher, J. Kelso, S. Paabo, and M. Mann. Deep proteome and transcriptome mapping of a human cancer cell line. Mol Syst Biol, 7:548, 2011. Nagaraj, Nagarjuna Wisniewski, Jacek R Geiger, Tamar Cox, Juergen Kircher, Martin Kelso, Janet Paabo, Svante Mann, Matthias Research Support, Non-U.S. Gov’t England Molecular systems biology Mol Syst Biol. 2011 Nov 8;7:548. doi: 10.1038/msb.2011.81.

44. A. Degeorges, F. Hoffschir, O. Cussenot, C. Gauville, A. Le Duc, B. Dutrillaux, and F. Calvo. Recurrent cytogenetic alterations of prostate carcinoma and amplification of c-myc or epidermal growth factor receptor in subclones of immortalized pnt1 human prostate epithelial cell line. Int J Cancer, 62(6):724–31, 1995. Degeorges, A Hoffschir, F Cussenot, O Gauville, C Le Duc, A Dutrillaux, B Calvo, F Research Support, Non-U.S. Gov’t United States International journal of cancer Int J Cancer. 1995 Sep 15;62(6):724–31.

45. M. Uhlen, E. Bjorling, C. Agaton, C. A. Szigyarto, B. Amini, E. Andersen, A. C. Andersson, P. Angelidou, A. Asplund, C. Asplund, L. Berglund, K. Bergstrom, H. Brumer, D. Cerjan, M. Ekstrom, A. Elobeid, C. Eriksson, L. Fagerberg, R. Falk, J. Fall, M. Forsberg, M. G. Bjorklund, K. Gumbel, A. Halimi, I. Hallin, C. Hamsten, M. Hansson, M. Hedhammar, G. Hercules, C. Kampf, K. Larsson, M. Lindskog, W. Lodewyckx, J. Lund, J. Lundeberg, K. Magnusson, E. Malm, P. Nilsson, J. Odling, P. Oksvold, I. Olsson, E. Oster, J. Ottosson, L. Paavilainen, A. Persson, R. Rimini, J. Rockberg, M. Runeson, A. Sivertsson, A. Skollermo, J. Steen, M. Stenvall, F. Sterky, S. Stromberg, M. Sundberg, H. Tegel, S. Tourle, E. Wahlund, A. Walden, J. Wan, H. Wernerus, J. Westberg, K. Wester, U. Wrethagen, L. L. Xu, S. Hober, and F. Ponten. A human protein atlas for normal and cancer tissues based on antibody proteomics. Mol Cell Proteomics, 4(12):1920–32, 2005. Uhlen, Mathias Bjorling, Erik Agaton, Charlotta Szigyarto, Cristina Al-Khalili Amini, Bahram Andersen, Elisabet Andersson, Ann-Catrin Angelidou, Pia Asplund, Anna Asplund, Caroline Berglund, Lisa Bergstrom, Kristina Brumer, Harry Cerjan, Dijana Ekstrom, Marica Elobeid, Adila Eriksson, Cecilia Fagerberg, Linn Falk, Ronny Fall, Jenny Forsberg, Mattias Bjorklund, Marcus Gry Gumbel, Kristoffer Halimi, Asif Hallin, Inga Hamsten, Carl Hansson, Marianne Hedhammar, My Hercules, Gorel Kampf, Caroline Larsson, Karin Lindskog, Mats Lodewyckx, Wald Lund, Jan Lundeberg, Joakim Magnusson, Kristina Malm, Erik Nilsson, Peter Odling, Jenny Oksvold, Per Olsson, Ingmarie Oster, Emma Ottosson, Jenny Paavilainen, Linda Persson, Anja Rimini, Rebecca Rockberg, Johan Runeson, Marcus Sivertsson, Asa Skollermo, Anna Steen, Johanna Stenvall, Maria Sterky, Fredrik Stromberg, Sara Sundberg, Marten Tegel, Hanna Tourle, Samuel Wahlund, Eva Walden, Annelie Wan, Jinghong Wernerus, Henrik Westberg, Joakim Wester, Kenneth Wrethagen, Ulla Xu, Lan Lan Hober, Sophia Ponten, Fredrik Research Support, Non-U.S. Gov’t United States Molecular & cellular proteomics: MCP Mol Cell Proteomics. 2005 Dec;4(12):1920–32. Epub 2005 Aug 27.

46. R. Gilad, K. Meir, I. Stein, L. German, E. Pikarsky, and N. J. Mabjeesh. High sept9_i1 protein expression is associated with high-grade prostate cancers. PLoS One, 10(4): e0124251, 2015. Gilad, Roni Meir, Karen Stein, Ilan German, Larissa Pikarsky, Eli Mabjeesh, Nicola J Research Support, Non-U.S. Gov’t United States PloS one PLoS One. 2015 Apr 21;10(4):e0124251. doi: 10.1371/journal.pone.0124251. eCollection 2015.

47. W. K. Leung, A. K. Ching, A. W. Chan, T. C. Poon, H. Mian, A. S. Wong, K. F. To, and N. Wong. A novel interplay between oncogenic pftk1 protein kinase and tumor suppressor tagln2 in the control of liver cancer cell motility. Oncogene, 30(44):4464–75, 2011. Leung, W K C Ching, A K K Chan, A W H Poon, T C W Mian, H Wong, A S T To, K-F Wong, N Research Support, Non-U.S. Gov’t England Oncogene Oncogene. 2011 Nov 3;30(44):4464–75. doi: 10.1038/onc.2011.161. Epub 2011 May 16.

48. K. Garber. Energy deregulation: licensing tumors to grow. Science, 312(5777):1158–9, 2006. Garber, Ken News United States Science (New York, N.Y.) Science. 2006 May 26;312(5777):1158–9.

49. K. R. Jakobsen, E. Sorensen, K. K. Brondum, T. F. Daugaard, R. Thomsen, and A. L. Nielsen. Direct rna sequencing mediated identification of mrna localized in protrusions of human mda-mb-231 metastatic breast cancer cells. J Mol Signal, 8(1):9, 2013. Jakobsen, Kristine Raaby Sorensen, Emilie Brondum, Karin Kathrine Daugaard,Tina Fuglsang Thomsen, Rune Nielsen, Anders Lade England Journal of molecular signaling J Mol Signal. 2013 Sep 1;8(1):9. doi: 10.1186/1750-2187-8-9.

50. R. Mayor and C. Carmona-Fontaine. Keeping in touch with contact inhibition of locomotion. Trends Cell Biol, 20(6):319–28, 2010. Mayor, Roberto Carmona-Fontaine, Carlos G0801145/Medical Research Council/United Kingdom Wellcome Trust/United Kingdom Medical Research Council/United Kingdom Biotechnology and Biological Sciences Research Council/United Kingdom Research Support, Non-U.S. Gov’t Review England Trends in cell biology Trends Cell Biol. 2010Jun;20(6):319–28. doi: 10.1016/j.tcb.2010.03.005.

51. A. S. Azmi, B. Bao, and F. H. Sarkar. Exosomes in cancer development, metastasis, and drug resistance: a comprehensive review. Cancer Metastasis Rev, 32(3-4):623–42, 2013. Azmi, Asfar S Bao, Bin Sarkar, Fazlul H R01 CA132794/CA/NCI NIH HHS/United States Review Netherlands Cancer metastasis reviews Cancer Metastasis Rev. 2013 Dec;32(3-4):623–42. doi: 10.1007/s10555-013-9441-9.

52. L. Rauschenberger, D. Staar, K. Thom, C. Scharf, S. Venz, G. Homuth, R. Schluter, L. O. Brandenburg, P. Ziegler, U. Zimmermann, W. Weitschies, U. Volker, U. Lendeckel, R. Walther, M. Burchardt, and M. B. Stope. Exosomal particles secreted by prostate cancer cells are potent mrna and protein vehicles for the interference of tumor and tumor environment. Prostate, 76(4):409–24, 2016. Rauschenberger, Lisa Staar, Doreen Thom, Kathleen Scharf, Christian Venz, Simone Homuth, Georg Schluter, Rabea Brandenburg, Lars-Ove Ziegler, Patrick Zimmermann, Uwe Weitschies, Werner Volker, Uwe Lendeckel, Uwe Walther, Reinhard Burchardt, Martin Stope, Matthias B United States The Prostate Prostate. 2016 Mar;76(4):409–24. doi: 10.1002/pros.23132. Epub 2015 Dec 8.

53. A. Y. Yam, Y. Xia, H. T Lin, A. Burlingame, M. Gerstein, and J. Frydman. Defining the tric/cct interactome links chaperonin function to stabilization of newly made proteins with complex topologies. Nat Struct Mol Biol, 15(12):1255–62, 2008. Yam, Alice Y Xia, Yu Lin, Hen-Tzu Jill Burlingame, Alma Gerstein, Mark Frydman, Judith R01 GM056433-11/GM/NIGMS NIH HHS/United States R01 GM056433-08/GM/NIGMS NIH HHS/United States R01 GM056433-06/GM/NIGMS NIH HHS/United States R01 GM056433/GM/NIGMS NIH HHS/United States R01 GM056433-07/GM/NIGMS NIH HHS/United States R01 GM056433-10/GM/NIGMS NIH HHS/United States R01 GM056433-10S1/GM/NIGMS NIH HHS/United States R01 GM056433-09/GM/NIGMS NIH HHS/United States Research Support, N.I.H., Extramural Research Support, Non-U.S. Gov’t United States Nature structural & molecular biology Nat Struct Mol Biol. 2008 Dec;15(12):1255–62. doi: 10.1038/nsmb.1515. Epub 2008 Nov 16.

54. S. H. Roh, M. Kasembeli, D. Bakthavatsalam, W. Chiu, and D. J. Tweardy. Contribution of the type ii chaperonin, tric/cct, to oncogenesis. Int J Mol Sci, 16(11): 26706–20, 2015. Roh, Soung-Hun Kasembeli, Moses Bakthavatsalam, Deenadayalan Chiu, Wah Tweardy, David J P41 GM103832/GM/NIGMS NIH HHS/United States PN2 EY016525/EY/NEI NIH HHS/United States P41GM103832/GM/NIGMS NIH HHS/United States PN2EY016525/EY/NEI NIH HHS/United States Research Support, N.I.H., Extramural Research Support, Non-U.S. Gov’t Review Switzerland International journal of molecular sciences Int J Mol Sci. 2015 Nov6;16(11):26706–20. doi: 10.3390/ijms161125975.

55. A. G. Trinidad, P. A. Muller, J. Cuellar, M. Klejnot, M. Nobis, J. M. Valpuesta, and K. H. Vousden. Interaction of p53 with the cct complex promotes protein folding and wild-type p53 activity. Mol Cell, 50(6):805–17,2013. Trinidad, Antonio Garcia Muller, Patricia A J Cuellar, Jorge Klejnot, Marta Nobis, Max Valpuesta, Jose Maria Vousden, Karen H 11-0626/Worldwide Cancer Research/United Kingdom Cancer Research UK/United Kingdom Research Support, Non-U.S. Gov’t United States Molecular cell Mol Cell. 2013 Jun 27;50(6):805–17. doi: 10.1016/j.molcel.2013.05.002. Epub 2013 Jun 6.

56. J. Bi, A. Huang, T. Liu, T. Zhang, and H. Ma. Expression of dna damage checkpoint 53bp1 is correlated with prognosis, cell proliferation and apoptosis in colorectal cancer. IntJ Clin Exp Pathol, 8(6):6070–82, 2015. Bi, Jianping Huang, Ai Liu, Tao Zhang, Tao Ma, Hong United States International journal of clinical and experimental pathology Int J Clin Exp Pathol. Jun 1;8(6):6070–82. eCollection 2015.

57. S. M. Harding, and R. G. Bristow. Discordance between phosphorylation and recruitment of 53bp1 in response to dna double-strand breaks. Cell Cycle, 11(7):1432–44, 2012. Harding, Shane M Bristow, Robert G Canadian Institutes of Health Research/Canada Research Support, Non-U.S. Gov’t United States Cell cycle (Georgetown, Tex.) Cell Cycle. 2012 Apr 1;11(7):1432–44. doi: 10.4161/cc.19824. Epub 2012 Apr 1.

58. K. Savitsky, Y. Ziv, A. Bar-Shira, S. Gilad, D. A. Tagle, S. Smith, T. Uziel, S. Sfez, J. Nahmias, A. Sartiel, R. L. Eddy, T. B. Shows, F. S. Collins, Y. Shiloh, and G. Rotman. A human gene (ddx10) encoding a putative dead-box rna helicase at 11q22-q23. Genomics, 33(2): 199–206, 1996. Savitsky, K Ziv, Y Bar-Shira, A Gilad, S Tagle, D A Smith, S Uziel, T Sfez, S Nahmias, J Sartiel, A Eddy, R L Shows, T B Collins, F S Shiloh, Y Rotman, G NS31763/NS/NINDS NIH HHS/United States Research Support, Non-U.S. Gov’t Research Support, U.S. Gov’t, Non-PH.S. Research Support, U.S. Gov’t, PH.S. United States Genomics Genomics. 1996 Apr 15;33(2):199–206.

59. L. Yang, C. Lin, and Z. R. Liu. Phosphorylations of dead box p68 rna helicase are associated with cancer development and cell proliferation. Mol Cancer Res, 3(6):355–63, 2005. Yang, Liuqing Lin, Chunru Liu, Zhi-Ren GM063874/GM/NIGMS NIH HHS/United States Research Support, N.I.H., Extramural Research Support, Non-U.S. Gov’t Research Support, U.S. Gov’t, PH.S. United States Molecular cancer research: MCR Mol Cancer Res. 2005 Jun;3(6):355–63.

60. E. A. Gustafson, and G. M. Wessel. Dead-box helicases: posttranslational regulation and function. Biochem Biophys Res Commun, 395(1):1–6, 2010. Gustafson, Eric A Wessel, Gary M R01 HD028152/HD/NICHD NIH HHS/United States R01 HD028152-16/HD/NICHD NIH HHS/United States T32 GM007601/GM/NIGMS NIH HHS/United States Review United States Biochemical and biophysical research communications Biochem Biophys Res Commun. 2010 Apr 23;395(1):1–6. doi: 10.1016/j.bbrc.2010.02.172. Epub 2010 Mar 3.

61. K. Lassi and N. A. Dawson. Update on castrate-resistant prostate cancer: 2010. Curr Opin Oncol, 22(3):263–7, 2010. Lassi, Kiran Dawson, Nancy A United States Current opinion in oncology Curr Opin Oncol. 2010 May;22(3):263–7. doi: 10.1097/CC0.0b013e3283380939.

62. M. V. Liberti, and J. W. Locasale. The warburg effect: How does it benefit cancer cells? Trends Biochem Sci, 41(3):211–8, 2016. Liberti, Maria V Locasale, Jason W R01 CA193256/CA/NCI NIH HHS/United States R01CA193256/CA/NCI NIH HHS/United States R00CA168997/CA/NCI NIH HHS/United States R00 CA168997/CA/NCI NIH HHS/United States T32 GM007273/GM/NIGMS NIH HHS/United States T32GM007273/GM/NIGMS NIH HHS/United States Research Support, N.I.H., Extramural Research Support, Non-U.S. Gov’t Research Support, U.S. Gov’t, Non-P.H.S. Review England Trends in biochemical sciences Trends Biochem Sci. 2016 Mar;41(3):211–8. doi: 10.1016/j.tibs.2015.12.001. Epub Jan 5.

63. J. P. Thiery, and J. P. Sleeman. Complex networks orchestrate epithelial-mesenchymal transitions. Nat Rev Mol Cell Biol, 7(2):131–42, 2006. Thiery, Jean Paul Sleeman, Jonathan P Research Support, Non-U.S. Gov’t Review England Nature reviews. Molecular cell biology Nat Rev Mol Cell Biol. 2006 Feb;7(2):131–42.

64. D. Hanahan and R. A. Weinberg. Hallmarks of cancer: the next generation. Cell, 144(5):646–74, 2011. Hanahan, Douglas Weinberg, Robert A Research Support, N.I.H., Extramural Review United States Cell Cell. 2011 Mar 4;144(5):646–74. doi: 10.1016/j.cell.2011.02.013.

65. C. S. Yang, T. A. Melhuish, A. Spencer, L. Ni, Y. Hao, K. Jividen, T. E. Harris, C. Snow, Jr. Frierson, H. F., D. Wotton, and B. M. Paschal. The protein kinase c super-family member pkn is regulated by mtor and influences differentiation during prostate cancer progression. Prostate, 77(15):1452–1467, 2017. Yang, Chun-Song Melhuish, Tiffany A Spencer, Adam Ni, Li Hao, Yi Jividen, Kasey Harris, Thurl E Snow, Chelsi Frierson, Henry F Jr Wotton, David Paschal, Bryce M P01 CA104106/CA/NCI NIH HHS/United States R01 DK101946/DK/NIDDK NIH HHS/United States R01 NS077958/NS/NINDS NIH HHS/United States T32 GM008136/GM/NIGMS NIH HHS/United States United States The Prostate Prostate. 2017 Nov;77(15):1452–1467. doi: 10.1002/pros.23400. Epub 2017 Sep 6.

66. K. Williams, R. Ghosh, P. V. Giridhar, G. Gu, T. Case, S. M. Belcher, and S. Kasper. Inhibition, of stathmin1 accelerates the metastatic process. Cancer Res, 72(20):5407–17, 2012. Williams, Karin Ghosh, Ritwik Giridhar, Premkumar Vummidi Gu, Guangyu Case, Thomas Belcher, Scott M Kasper, Susan R01 DK059142/DK/NIDDK NIH HHS/United States R01 DK060957/DK/NIDDK NIH HHS/United States R01 DK60957/DK/NIDDK NIH HHS/United States Research Support, N.I.H., Extramural Research Support, Non-U.S. Gov’t Research Support, U.S. Gov’t, Non-PH.S. United States Cancer research Cancer Res. 2012 Oct 15;72(20):5407–17. doi: 10.1158/0008-5472.CAN-12-1158. Epub 2012 Aug 21.

67. S. W. Plouffe, Z. Meng, K. C. Lin, B. Lin, A. W. Hong, J. V. Chun, and K. L. Guan. Characterization of hippo pathway components by gene inactivation. Mol Cell, 64(5):993–1008, 2016. Plouffe, Steven W Meng, Zhipeng Lin, Kimberly C Lin, Brian Hong, Audrey W Chun, Justin V Guan, Kun-Liang R01 DE015964/DE/NIDCR NIH HHS/United States R01 GM051586/GM/NIGMS NIH HHS/United States R35 CA196878/CA/NCI NIH HHS/United States T32 GM007752/GM/NIGMS NIH HHS/United States United States Molecular cell Mol Cell. 2016 Dec 1;64(5):993–1008. doi: 10.1016/j.molcel.2016.10.034.

68. Jong-Moon Park, Ji-Hwan Park, Dong-Gi Mun, Jingi Bae, Jae Hun Jung, Seunghoon Back, Hangyeore Lee, Hokeun Kim, Hee-Jung Jung, Hark Kyun Kim, Hookeun Lee, Kwang Pyo Kim, Daehee Hwang, and Sang-Won Lee. Integrated analysis of global proteome, phosphoproteome, and glycoproteome enables complementary interpretation of disease-related protein networks. Scientific Reports, 5:18189, 2015.

69. H. Tan, K. Yang, Y. Li, T. I. Shaw, Y. Wang, D. B. Blanco, X. Wang, J. H. Cho, H. Wang, S. Rankin, C. Guy, J. Peng, and H. Chi. Integrative proteomics and phosphoproteomics profiling reveals dynamic signaling networks and bioenergetics pathways underlying t cell activation. Immunity, 46(3):488–503, 2017. Tan, Haiyan Yang, Kai Li, Yuxin Shaw, Timothy I Wang, Yanyan Blanco, Daniel Bastardo Wang, Xusheng Cho, Ji-Hoon Wang, Hong Rankin, Sherri Guy, Cliff Peng, Junmin Chi, Hongbo R01 AG053987/AG/NIA NIH HHS/United States R01 GM114260/GM/NIGMS NIH HHS/United States R01 NS064599/NS/NINDS NIH HHS/United States P30 CA021765/CA/NCI NIH HHS/United States R01 AI105887/AI/NIAID NIH HHS/United States R01 CA176624/CA/NCI NIH HHS/United States R01 AI101407/AI/NIAID NIH HHS/United States R01 AG047928/AG/NIA NIH HHS/United States United States Immunity Immunity. Mar 21;46(3):488–503. doi: 10.1016/j.immuni.2017.02.010. Epub 2017 Mar 9.

70. K. Katada, T. Tomonaga, M. Satoh, K. Matsushita, Y. Tonoike, Y. Kodera, T. Hanazawa, F. Nomura, and Y. Okamoto. Plectin promotes migration and invasion of cancer cells and is a novel prognostic marker for head and neck squamous cell carcinoma. J Proteomics, 75 (6):1803–15, 2012. Katada, Koji Tomonaga, Takeshi Satoh, Mamoru Matsushita, Kazuyuki Tonoike, Yurie Kodera, Yoshio Hanazawa, Toyoyuki Nomura, Fumio Okamoto, Yoshitaka Research Support, Non-U.S. Gov’t Netherlands Journal of proteomics J Proteomics. 2012 Mar 16;75(6):1803–15. doi: 10.1016/j.jprot.2011.12.018. Epub 2011 Dec 30.

71. M. Sutoh Yoneyama, S. Hatakeyama, T. Habuchi, T. Inoue, T. Nakamura, T. Funyu,G. Wiche, C. Ohyama, and S. Tsuboi. Vimentin intermediate filament and plectin provide a scaffold for invadopodia, facilitating cancer cell invasion and extravasation for metastasis. Eur J Cell Biol, 93(4):157–69, 2014. Sutoh Yoneyama, Mihoko Hatakeyama, Shingo Habuchi, Tomonori Inoue, Takamitsu Nakamura, Toshiya Funyu, Tomihisa Wiche, Gerhard Ohyama, Chikara Tsuboi, Shigeru Research Support, Non-U.S. Gov’t Germany European journal of cell biology Eur J Cell Biol. 2014 Apr;93(4):157–69. doi: 10.1016/j.ejcb.2014.03.002. Epub 2014 Apr 15.

72. T. C. Burch, M. T. Watson, and J. O. Nyalwidhe. Variable metastatic potentials correlate with differential plectin and vimentin expression in syngeneic androgen independent prostate cancer cells. PLoS One, 8(5):e65005, 2013. Burch, Tanya C Watson, Megan T Nyalwidhe, Julius O Research Support, Non-U.S. Gov’t United States PloS one PLoS One. 2013 May 22;8(5):e65005. doi: 10.1371/journal.pone.0065005. Print 2013.

73. Y Luo, F. Kong, Z. Wang, D. Chen, Q. Liu, T. Wang, R. Xu, X. Wang, and J. Y Yang. Loss of asap3 destabilizes cytoskeletal protein actg1 to suppress cancer cell migration. Mol Med Rep, 9(2):387–94, 2014. Luo, Yu Kong, Fang Wang, Zhen Chen, Dahan Liu, Qiuyan Wang, Tao Xu, Ruian Wang, Xianyuan Yang, James Y Research Support, Non-U.S. Gov’t Greece Molecular medicine reports Mol Med Rep. 2014 Feb;9(2):387–94. doi: 10.3892/mmr.2013.1831. Epub 2013 Nov 27.

74. A. Bretscher, K. Edwards, and R. G. Fehon. Erm proteins and merlin: integrators at the cell cortex. Nat Rev Mol Cell Biol, 3(8):586–99, 2002. Bretscher, Anthony Edwards, Kevin Fehon, Richard G R01 NS034783/NS/NINDS NIH HHS/United States Research Support, U.S. Gov’t, Non-P.H.S. Research Support, U.S. Gov’t, P.H.S. Review England Nature reviews. Molecular cell biology Nat Rev Mol Cell Biol. 2002 Aug;3(8):586–99. doi: 10.1038/nrm882.

75. Y. Saygideger-Kont, T. Z. Minas, H. Jones, S. Hour, H. Celik, I. Temel, J. Han, N. Atabey, H. V. Erkizan, J. A. Toretsky, and A. Uren. Ezrin enhances egfr signaling and modulates erlotinib sensitivity in non-small cell lung cancer cells. Neoplasia, 18(2):111–20, 2016. Saygideger-Kont, Yasemin Minas, Tsion Zewdu Jones, Hayden Hour, Sarah Celik, Haydar Temel, Idil Han, Jenny Atabey, Nese Erkizan, Hayriye Verda Toretsky, Jeffrey A Uren, Aykut P30 CA051008/CA/NCI NIH HHS/United States P30 CA51008/CA/NCI NIH HHS/United States Research Support, N.I.H., Extramural United States Neoplasia (New York, N.Y.) Neoplasia. 2016 Feb;18(2):111–20. doi: 10.1016/j.neo.2016.01.002.

76. D. Soave, H. Corvol, N. Panjwani, J. Gong, W. Li, P. Y. Boelle, P. R. Durie, A. D. Paterson, J. M. Rommens, L. J. Strug, and L. Sun. A joint location-scale test improves power to detect associated snps, gene sets, and pathways. Am J Hum Genet, 97(1): 125–38, 2015. Soave, David Corvol, Harriet Panjwani, Naim Gong, Jiafen Li, Weili Boelle, Pierre-Yves Durie, Peter R Paterson, Andrew D Rommens, Johanna M Strug, Lisa J Sun, Lei 201309MOP-310732-G-CEAA-117978/Canadian Institutes of Health Research/Canada MOP-258916/Canadian Institutes of Health Research/Canada Evaluation Studies Research Support, Non-U.S. Gov’t United States American journal of human genetics Am J Hum Genet. 2015 Jul 2;97(1):125–38. doi: 10.1016/j.ajhg.2015.05.015.

77. X. Y. Kuang, H. S. Jiang, K. Li, Y. Z. Zheng, Y. R. Liu, F. Qiao, S. Li, X. Hu, and Z. M. Shao. The phosphorylation-specific association of stmn1 with grp78 promotes breast cancer metastasis. Cancer Lett, 377(1):87–96, 2016. Kuang, Xia-Ying Jiang, He-Sheng Li, Kai Zheng, Yi-Zi Liu, Yi-Rong Qiao, Feng Li, Shan Hu, Xin Shao, Zhi-Ming Research Support, Non-U.S. Gov’t Ireland Cancer letters Cancer Lett. 2016 Jul 10;377(1):87–96. doi: 10.1016/j.canlet.2016.04.035. Epub 2016 Apr 26.

78. S. Germann, L. Gratadou, M. Dutertre, and D. Auboeuf. Splicing programs and cancer. J Nucleic Acids, 2012:269570, 2012. Germann, Sophie Gratadou, Lise Dutertre, Martin Auboeuf, Didier United States Journal of nucleic acids J Nucleic Acids. 2012;2012:269570. Epub 2011 Oct 24.

79. J. Munkley, K. Livermore, P. Rajan, and D. J. Elliott. Rna splicing and splicing regulator changes in prostate cancer pathology. Hum Genet, 2017. Munkley, Jennifer Livermore, Karen Rajan, Prabhakar Elliott, David J Review Germany Human genetics Hum Genet. 2017 Apr 5. doi: 10.1007/s00439-017-1792-9.

80. J. Chen, J. Zhou, C. K. Sanders, J. P. Nolan, and H. Cai. A surface display yeast two-hybrid screening system for high-throughput protein interactome mapping. Anal Biochem, 390 (1):29–37, 2009. Chen, Jun Zhou, Jianhong Sanders, Claire K Nolan, John P Cai, Hong EB003824/EB/NIBIB NIH HHS/United States RR01315/RR/NCRR NIH HHS/United States Research Support, N.I.H., Extramural United States Analytical biochemistry Anal Biochem. 2009 Jul 1;390(1):29–37. Epub 2009 Mar 17.

81. C. Naro and C. Sette. Phosphorylation-mediated regulation of alternative splicing in cancer. Int J Cell Biol, 2013:151839, 2013. Naro, Chiara Sette, Claudio Review International journal of cell biology Int J Cell Biol. 2013;2013:151839. Epub 2013 Aug 28.

82. H. Molina, D. M. Horn, N. Tang, S. Mathivanan, and A. Pandey. Global proteomic profiling of phosphopeptides using electron transfer dissociation tandem mass spectrometry. Proc Natl Acad Sci U S A, 104(7):2199–204, 2007. Molina, Henrik Horn, David M Tang, Ning Mathivanan, Suresh Pandey, Akhilesh R01CA106424/CA/NCI NIH HHS/United States U54 RR020839/RR/NCRR NIH HHS/United States N01-HV-28180/HV/NHLBI NIH HHS/United States R01 CA106424/CA/NCI NIH HHS/United States U54RR020839/RR/NCRR NIH HHS/United States N01HV28180/HL/NHLBI NIH HHS/United States Research Support, N.I.H., Extramural Research Support, Non-U.S. Gov’t United States Proceedings of the National Academy of Sciences of the United States of America Proc Natl Acad Sci U S A. 2007 Feb 13;104(7):2199–204. Epub 2007 Feb 7.

83. J. E. Mermoud, P. Cohen, and A. I. Lamond. Ser/thr-specific protein phosphatases are required for both catalytic steps of pre-mrna splicing. Nucleic Acids Res, 20(20):5263–9, 1992. Mermoud, J E Cohen, P Lamond, A I Research Support, Non-U.S. Gov’t England Nucleic acids research Nucleic Acids Res. 1992 Oct 25;20(20):5263–9.

84. S. Broderick, K. Rehmet, C. Concannon, and H. P. Nasheuer. Eukaryotic single-stranded dna binding proteins: central factors in genome stability. Subcell Biochem, 50:143–63, 2010. Broderick, Sandra Rehmet, Kristina Concannon, Claire Nasheuer, Heinz-Peter Research Support, Non-U.S. Gov’t Review United States Sub-cellular biochemistry Subcell Biochem. 2010;50:143–63. doi: 10.1007/978-90-481-3471-7_8.

85. M. Dhawan, C. J. Ryan, and A. Ashworth. Dna repair deficiency is common in advanced prostate cancer: New therapeutic opportunities. Oncologist, 21(8):940–5, 2016. Dhawan, Mallika Ryan, Charles J Ashworth, Alan Review United States The oncologist Oncologist. 2016 Aug;21(8):940–5. doi: 10.1634/theoncologist.2016-0135. Epub 2016 Jun 17.

86. A. Montecucco and G. Biamonti. Pre-mrna processing factors meet the dna damage response. Front Genet, 4:102, 2013. Montecucco, Alessandra Biamonti, Giuseppe Switzerland Frontiers in genetics Front Genet. 2013 Jun 6;4:102. doi: 10.3389/fgene.2013.00102. eCollection 2013.

87. X. Jacq, M. Kemp, N. M. Martin, and S. P. Jackson. Deubiquitylating enzymes and dna damage response pathways. Cell Biochem Biophys, 67(1):25–43, 2013. Jacq, Xavier Kemp, Mark Martin, Niall M B Jackson, Stephen P 092096/Wellcome Trust/United Kingdom 11224/Cancer Research UK/United Kingdom 268536/European Research Council/International A11224/Cancer Research UK/United Kingdom Research Support, Non-U.S. Gov’t Review United States Cell biochemistry and biophysics Cell Biochem Biophys. 2013 Sep;67(1):25–43. doi: 10.1007/s12013-013-9635-3.

88. F. Yuan, G. Li, and T. Tong. Nucleolar and coiled-body phosphoprotein 1 (nolc1) regulates the nucleolar retention of trf2. Cell Death Discov, 3:17043, 2017. Yuan, Fuwen Li, Guodong Tong, Tanjun United States Cell death discovery Cell Death Discov. 2017 Sep 4;3:17043. doi: 10.1038/cddiscovery.2017.43. eCollection 2017.

89. R. Fagerlund, L. Kinnunen, M. Kohler, I. Julkunen, and K. Melen. Nf-kappab is transported into the nucleus by importin alpha3 and importin alpha4. J Biol Chem, 280(16):15942–51, 2005. Fagerlund, Riku Kinnunen, Leena Kohler, Matthias Julkunen, Ilkka Melen, Krister Research Support, Non-U.S. Gov’t United States The Journal of biological chemistry J Biol Chem. 2005 Apr 22;280(16):15942–51. doi: 10.1074/jbc.M500814200. Epub 2005 Jan 27.

90. R. J. Jin, Y. Lho, L. Connelly, Y. Wang, X. Yu, L. Saint Jean, T. C. Case, K. Ellwood Yen, C. L. Sawyers, N. A. Bhowmick, T. S. Blackwell, F. E. Yull, and R. J. Matusik. The nuclear factor-kappab pathway controls the progression of prostate cancer to androgen-independent growth. Cancer Res, 68(16):6762–9, 2008. Jin, Ren Jie Lho, Yongsoo Connelly, Linda Wang, Yongqing Yu, Xiuping Saint Jean, Leshana Case, Thomas C Ellwood-Yen, Katharine Sawyers, Charles L Bhowmick, Neil A Blackwell, Timothy S Yull, Fiona E Matusik, Robert J R01-HL061419/HL/NHLBI NIH HHS/United States R01 AG023490-04/AG/NIA NIH HHS/United States R01 HL061419-08/HL/NHLBI NIH HHS/United States R01 CA076142/CA/NCI NIH HHS/United States R01 HL061419/HL/NHLBI NIH HHS/United States R01 CA076142-11A1/CA/NCI NIH HHS/United States R01 AG023490/AG/NIA NIH HHS/United States R01-AG023490/AG/NIA NIH HHS/United States R01-CA76142/CA/NCI NIH HHS/United States Research Support, N.I.H., Extramural Research Support, Non-U.S. Gov’t United States Cancer research Cancer Res. 2008 Aug 15;68(16):6762–9. doi: 10.1158/0008-5472.CAN-08-0107.

91. A. Y. Lai and P. A. Wade. Cancer biology and nurd: a multifaceted chromatin remodelling complex. Nat Rev Cancer, 11(8):588–96, 2011. Lai, Anne Y Wade, Paul A Z01 ES101965/ES/NIEHS NIH HHS/United States ZIA ES101965-07/Intramural NIH HHS/United States Z01ES101965/ES/NIEHS NIH HHS/United States Research Support, N.I.H., Extramural Research Support, N.I.H., Intramural Review England Nature reviews. Cancer Nat Rev Cancer. 2011 Jul 7;11(8):588–96. doi: 10.1038/nrc3091.

92. C. G. Spruijt, M. S. Luijsterburg, R. Menafra, R. G. Lindeboom, P. W. Jansen, R. R. Edupuganti, M. P. Baltissen, W. W. Wiegant, M. C. Voelker-Albert, F. Matarese, A. Mensinga, I. Poser, H. R. Vos, H. G. Stunnenberg, H. van Attikum, and M. Vermeulen. Zmynd8 colocalizes with nurd on target genes and regulates poly(adp-ribose)-dependent recruitment of gatad2a/nurd to sites of dna damage. Cell Rep, 17(3):783–798, 2016. Spruijt, Cornelia G Luijsterburg, Martijn S Menafra, Roberta Lindeboom, Rik G H Jansen, Pascal W T C Edupuganti, Raghu Ram Baltissen, Marijke P Wiegant, Wouter W Voelker-Albert, Moritz C Matarese, Filomena Mensinga, Anneloes Poser, Ina Vos, Harmjan R Stunnenberg, Hendrik G van Attikum, Haico Vermeulen, Michiel United States Cell reports Cell Rep. 2016 Oct 11;17(3):783–798. doi: 10.1016/j.celrep.2016.09.037.

93. S. Heeboll, M. Borre, P. D. Ottosen, C. L. Andersen, F. Mansilla, L. Dyrskjot, T. F. Orntoft, and N. Torring. Smarcc1 expression is upregulated in prostate cancer and positively correlated with tumour recurrence and dedifferentiation. Histol Histopathol, 23(9):1069–76, 2008. Heeboll, S Borre, M Ottosen, P D Andersen, C L Mansilla, F Dyrskjot, L Orntoft, T F Torring, N Research Support, Non-U.S. Gov’t Spain Histology and histopathology Histol Histopathol. 2008 Sep;23(9):1069–76. doi: 10.14670/HH-23.1069.

94. R. Hu, C. Lu, E. A. Mostaghel, S. Yegnasubramanian, M. Gurel, C. Tannahill, J. Edwards, W. B. Isaacs, P. S. Nelson, E. Bluemn, S. R. Plymate, and J. Luo. Distinct transcriptional programs mediated by the ligand-dependent full-length androgen receptor and its splice variants in castration-resistant prostate cancer. Cancer Res, 72(14):3457–62, 2012. Hu, Rong Lu, Changxue Mostaghel, Elahe A Yegnasubramanian, Srinivasan Gurel, Meltem Tannahill, Clare Edwards, Joanne Isaacs, William B Nelson, Peter S Bluemn, Eric Plymate, Stephen R Luo, Jun P50CA58286/CA/NCI NIH HHS/United States P50 CA058236/CA/NCI NIH HHS/United States P50 CA097186/CA/NCI NIH HHS/United States P50 CA97186/CA/NCI NIH HHS/United States P01 CA085859/CA/NCI NIH HHS/United States Research Support, N.I.H., Extramural Research Support, Non-U.S. Gov’t Research Support, U.S. Gov’t, Non-P.H.S. United States Cancer research Cancer Res. 2012 Jul 15;72(14):3457–62. doi: 10.1158/0008-5472.CAN-11-3892. Epub 2012 Jun 18.

95. Q. Wang, W. Li, Y. Zhang, X. Yuan, K. Xu, J. Yu, Z. Chen, R. Beroukhim, H. Wang, M. Lupien, T. Wu, M. M. Regan, C. A. Meyer, J. S. Carroll, A. K. Manrai, O. A. Janne, S. P. Balk, R. Mehra, B. Han, A. M. Chinnaiyan, M. A. Rubin, L. True, M. Fiorentino, C. Fiore, M. Loda, P. W. Kantoff, X. S. Liu, and M. Brown. Androgen receptor regulates a distinct transcription program in androgen-independent prostate cancer. Cell, 138(2): 245–56, 2009. Wang, Qianben Li, Wei Zhang, Yong Yuan, Xin Xu, Kexin Yu, Jindan Chen, Zhong Beroukhim, Rameen Wang, Hongyun Lupien, Mathieu Wu, Tao Regan, Meredith M Meyer, Clifford A Carroll, Jason S Manrai, Arjun Kumar Janne, Olli A Balk, Steven P Mehra, Rohit Han, Bo Chinnaiyan, Arul M Rubin, Mark A True, Lawrence Fiorentino, Michelangelo Fiore, Christopher Loda, Massimo Kantoff, Philip W Liu, X Shirley Brown, Myles P50 CA090381/CA/NCI NIH HHS/United States P50 CA090381-060005/CA/NCI NIH HHS/United States K99 CA126160/CA/NCI NIH HHS/United States P50CA090381/CA/NCI NIH HHS/United States K99 CA129565/CA/NCI NIH HHS/United States P50 CA090381-070005/CA/NCI NIH HHS/United States K99 CA129565-01A1/CA/NCI NIH HHS/United States Research Support, N.I.H., Extramural Research Support, U.S. Gov’t, Non-P.H.S. United States Cell Cell. 2009 Jul 23;138(2):245–56. doi: 10.1016/j.cell.2009.04.056.

96. E. J. Faivre, D. Wilcox, X. Lin, P. Hessler, M. Torrent, W. He, T. Uziel, D. H. Albert, K. McDaniel, W. Kati, and Y. Shen. Exploitation of castration-resistant prostate cancer transcription factor dependencies by the novel bet inhibitor abbv-075. Mol Cancer Res, 15(1):35–44, 2017. Faivre, Emily J Wilcox, Denise Lin, Xiaoyu Hessler, Paul Torrent, Maricel He, Wei Uziel, Tamar Albert, Daniel H McDaniel, Keith Kati, Warren Shen, Yu United States Molecular cancer research: MCR Mol Cancer Res. 2017 Jan;15(1):35–44. doi: 10.1158/1541-7786.MCR-16-0221. Epub 2016 Oct 5.

97. K. J. Pienta, C. Abate-Shen, D. B. Agus, R. M. Attar, L. W. Chung, N. M. Green berg, W. C. Hahn, J. T. Isaacs, N. M. Navone, D. M. Peehl, J. W. Simons, D. B. Solit, H. R. Soule, T. A. VanDyke, M. J. Weber, L. Wu, and R. L. Vessella. The current state of preclinical prostate cancer animal models. Prostate, 68(6):629–39, 2008. Pienta, Kenneth J Abate-Shen, Cory Agus, David B Attar, Ricardo M Chung, Leland W K Greenberg, Norman M Hahn, William C Isaacs, John T Navone, Nora M Peehl, Donna M Simons, Jonathon W Solit, David B Soule, Howard R VanDyke, Terry A Weber, Michael J Wu, Lily Vessella, Robert L U01 CA84296/CA/NCI NIH HHS/United States R01 CA105402/CA/NCI NIH HHS/United States R01 CA105402-01/CA/NCI NIH HHS/United States U19 CA113317/CA/NCI NIH HHS/United States U19 CA113317-01/CA/NCI NIH HHS/United States U01 CA084294/CA/NCI NIH HHS/United States P01 CA098912-01A1/CA/NCI NIH HHS/United States U01 CA084296/CA/NCI NIH HHS/United States P01CA098912/CA/NCI NIH HHS/United States P01 CA098912/CA/NCI NIH HHS/United States P01CA093900/CA/NCI NIH HHS/United States P01 CA10410605/CA/NCI NIH HHS/United States 2P50 CA69568/CA/NCI NIH HHS/United States R01 CA101904/CA/NCI NIH HHS/United States U01 CA084296-01/CA/NCI NIH HHS/United States R01 CA101904-01/CA/NCI NIH HHS/United States P50 CA069568-05/CA/NCI NIH HHS/United States P50 CA069568/CA/NCI NIH HHS/United States P01 CA104106/CA/NCI NIH HHS/United States P01 CA093900/CA/NCI NIH HHS/United States P01 CA093900-01A2/CA/NCI NIH HHS/United States Multicenter Study Research Support, N.I.H., Extramural Research Support, Non-U.S. Gov’t Research Support, U.S. Gov’t, Non-P.H.S. United States The Prostate Prostate. 2008 May 1;68(6):629–39. doi: 10.1002/pros.20726.

98. D. Cunningham and Z. You. In vitro and in vivo model systems used in prostate cancer research. J Biol Methods, 2(1), 2015. Cunningham, David You, Zong bing P20 GM103518/GM/NIGMS NIH HHS/United States P20 RR020152/RR/NCRR NIH HHS/United States R01 CA174714/CA/NCI NIH HHS/United States United States Journal of biological methods J Biol Methods. 2015;2(1). doi: 10.14440/jbm.2015.63.

99. F. Ardito, M. Giuliani, D. Perrone, G. Troiano, and L. Lo Muzio. The crucial role of protein phosphorylation in cell signaling and its use as targeted therapy (review). Int J Mol Med, 40 (2):271–280, 2017. Ardito, Fatima Giuliani, Michele Perrone, Donatella Troiano, Giuseppe Lo Muzio, Lorenzo Review Greece International journal of molecular medicine Int J Mol Med. 2017 Aug;40(2):271–280. doi: 10.3892/ijmm.2017.3036. Epub 2017 Jun 22.

