## Supplementary Information for "Integrative proteomic and phosphoproteomic profiling of prostate cell lines"

### Supplementary Tables

- **Supplementary Table 1:** proteins identified in the MS experiment, and subset of filtered proteins associated with at least 2 valid quantification values in all four cell lines, which were kept for expression analyses.
- **Supplementary Table 2:** phosphosites identified in the MS experiment, and subset of filtered phosphosites associated with at least 2 valid quantification values in all four cell lines, which were kept for expression analyses.
- **Supplementary Table 3:** subdatasets of interest in proteomic expression analyses. It contains the ANOVA-significant proteins, the proteins up- and downregulated in the three Prostate Cancer (PCa) cell lines as compared to the benign PNT1A cell line, the proteins up- and downregulated in the castration-resistant (CR: DU145 and PC3) cell lines as compared to the castration-sensitive (CS: LNCaP) cell line, and the proteins identified only in the CR or CS contexts (CR_only, CS_only).
- **Supplementary Table 4:** subdatasets of interest in phosphoproteomic expression analyses. It contains the ANOVA-significant phosphosites, the phosphosites up- and downregulated in the three Prostate Cancer (PCa) cell lines as compared to the benign PNT1A cell line, the phosphosites up- and downregulated in the castration-resistant (CR: DU145 and PC3) cell lines as compared to the castration-sensitive (CS: LNCaP) cell line, and the phosphosites identified only in the CR or CS contexts (CR_only, CS_only). It further contains the results of the KSEA analysis.
- **Supplementary Table 5:** raw results of the functional enrichment analyses with G:profiler and Ingenuity Pathway Analyses (IPA) .

### Supplementary Figures

- **Supplementary Figure 1:** dynamic range of the prostate cancer proteome. (a) Ranking of the absolute abundance using the IBAQ intensity. The expression values of every protein in the three replicates of the four studied cell lines were considered. (b) Zoom on the left box in (a) displaying the 25 less abundant proteins. (c) Zoom on the right box in (a) displaying the 25 most abundant proteins.
- **Supplementary Figure 2:** correlation between proteomic and phosphoproteomic expression values. We computed for each cell line the correlation between the expression values of the 135 proteins that were quantified both at the proteomic and the phosphoproteomic levels. For proteomics data, we computed the mean of the three replicated. For phosphoproteomics data, we computed the mean for all the phosphosites belonging to the same protein.
- **Supplementary Figure 3:** expression Profiles associated with Septin-9 (SEPT9). (a) Boxplot showing the SEPT9 protein expression values in the four cell lines under study. (b) Boxplot revealing the SEPT9 Serine-30 phosphosite expression values in the four cell lines under study.
