## Supplementary figures and images for "Integrative proteomic and phosphoproteomic profiling of prostate cell lines"

### Supplementary Figure 1

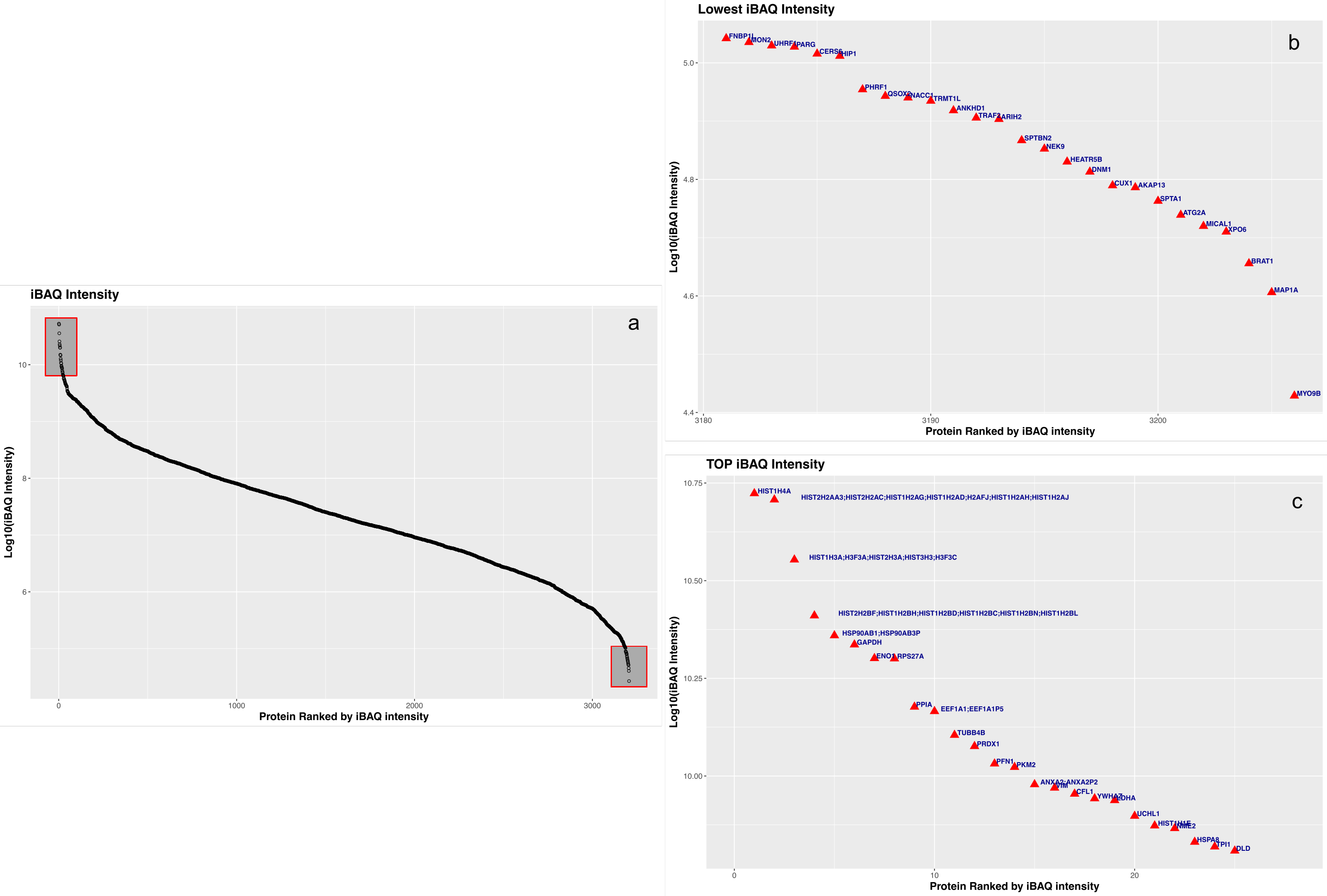

### Supplementary Figure 2

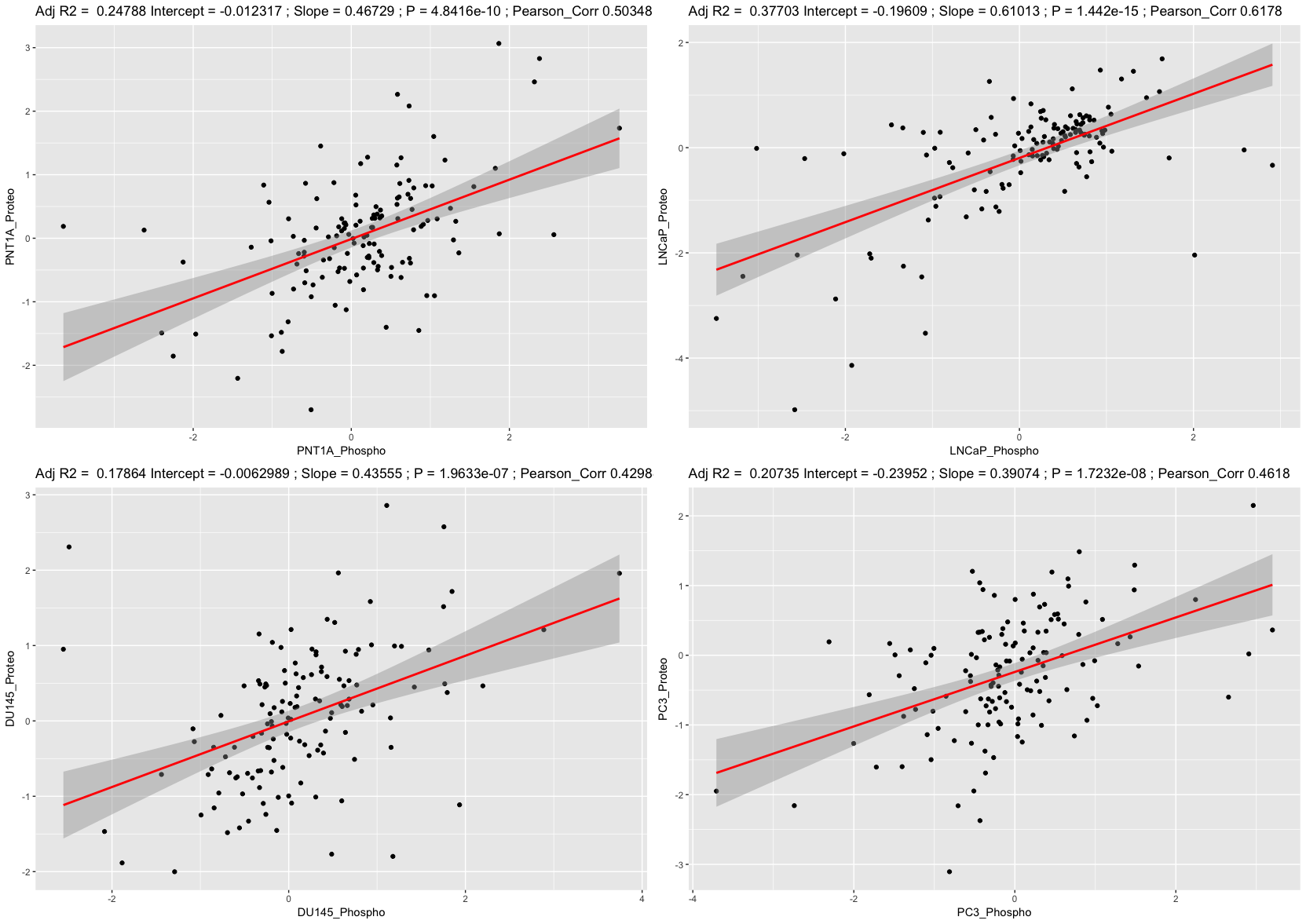

### Supplementary Figure 3

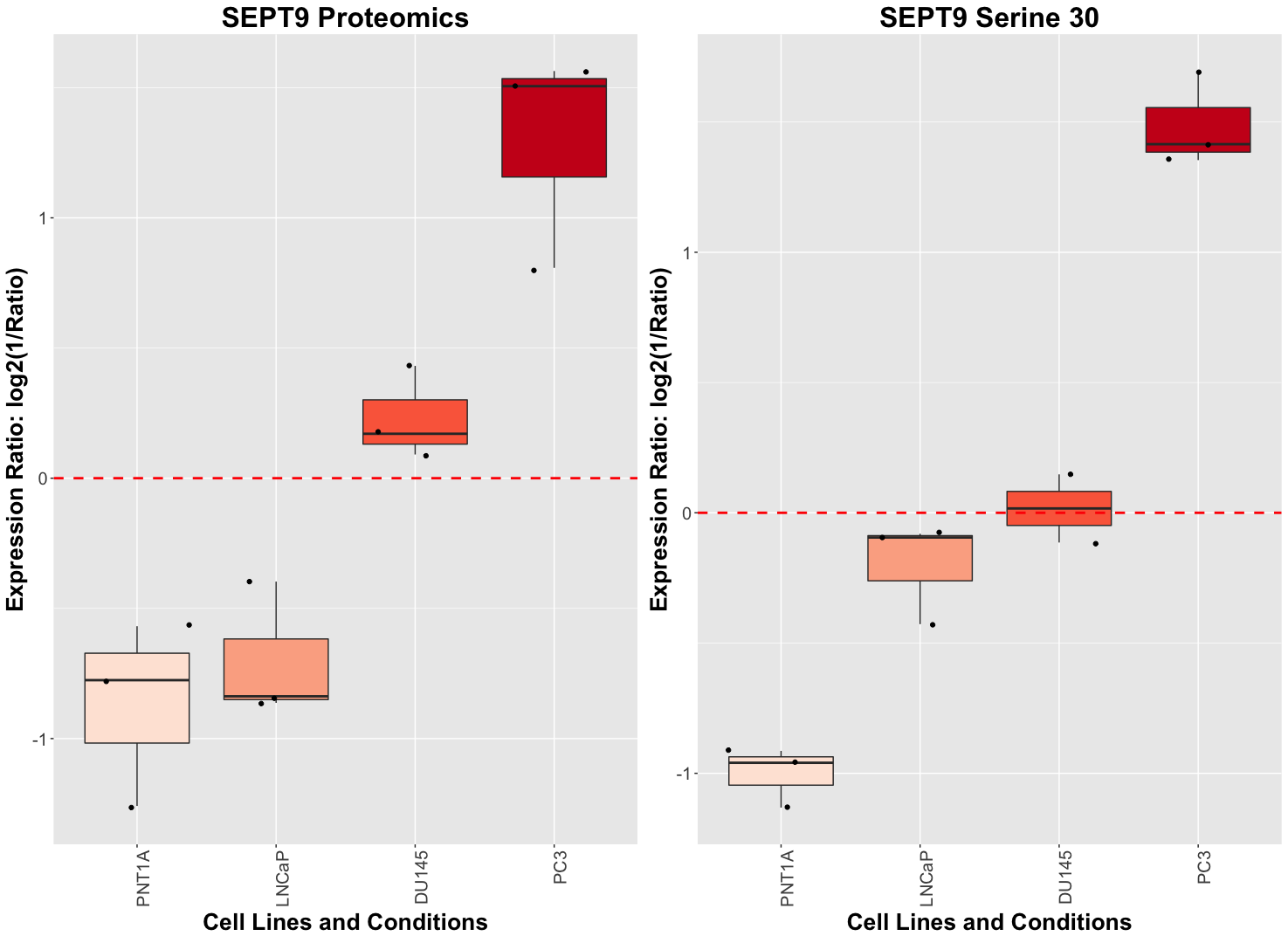
